# Somatic structural variation discovery on cancer pangenome graphs

**DOI:** 10.64898/2026.11.05.764632

**Authors:** Muti Kara, Akmuhammet Ashyralyyev, Can Alkan

## Abstract

Somatic structural variants (SVs) drive tumorigenesis across most cancers, yet low variant allele frequencies in subclonal populations often necessitate multi-sample sequencing per patient. Existing somatic SV callers rely on whole-genome read alignments against a single linear reference, incurring high computational costs and reference bias. We present janus, which discovers somatic SVs directly within a personal pangenome, a colored de Bruijn-like graph of locally consistent parsing (LCP) cores built from matched normal and tumor PacBio HiFi reads. janus identifies candidate SV sites traversed exclusively by tumor reads without prior alignment, mapping only short read snippets surrounding each site. Across five tumor-normal pairs in the CASTLE panel, jancs leads the 18-caller consensus with a mean Fl score of 0.887 (next-best caller: 0.858), widening its lead under low-signal regimes (0.489 versus 0.362 at 5×; 0.683 versus 0.520 at 20% purity). End to end, janus analyzes a tumor-normal pair in ~11 minutes on average with 16 threads, which is ~24× faster than read alignment alone on the same machine. In a 20-tumor patient simulation, joint graph calling outperforms independent pairwise runs in speed and accuracy, particularly at low depth. By processing multi-sample co-horts jointly and maintaining sensitivity at low coverage and purity, janus provides an efficient, reference-unbiased framework for cancer genomics.

jancs is available at https://github.com/BilkentCompGen/janus.

## 1 Introduction

Structural variants (SVs) are genomic alterations that span at least 50 base pairs (bp) [1]. In addition to inherited variants, cancer cells acquire SVs through flawed DNA replication and repair, from small deletions to rearrangements of entire chromosomes [2, 3]. SVs represent the primary class of driver mutations across most tumor types [2], and the resulting genomic instability fuels the acquisition of cancer hallmarks [4, 5]. Because cell lineages inherit these alterations [6], ongoing structural variation drives clonal diversification and tumor progression [7]. Consequently, rapid and accurate somatic SV identification is critical for clinical oncology [8, 9], yet remains technically demanding [10].

Early approaches relied on short reads to infer SVs indirectly from discordant read pairs, split reads, depth fluctuations, and local assembly [11, 12, 13]. However, short reads map poorly to repetitive regions, limiting detection accuracy [1]. Long-read sequencing from Oxford Nanopore Technologies (ONT) and PacBio HiFi overcomes mapping ambiguity because individual reads can span repetitive elements or entire SVs [14, 15, 16]. Integrating long reads resolved this limitation, uncovering 3 to 7 times as many SVs per genome as short-read datasets alone [1]. Between the two platforms, PacBio HiFi reads achieve 99.8% average base accuracy [17], making them better suited than ONT reads for exact sequence matching and breakpoint resolution despite their shorter lengths [15, 18].

Most studies analyze somatic SVs in bulk samples that pool DNA from thousands of cells [19]. Because ongoing mutations create distinct subclones [7], many variants occur only in small cell subpopulations. Normal-cell contamination dilutes these mutations that often fall below standard variant allele frequency (VAF) thresholds [9, 7]. Single-cell methods directly isolate distinct clones [20], but they introduce coverage gaps and allelic dropout that impede variant discovery [21]. Bulk sequencing therefore remains the standard approach. To improve clonal resolution, researchers sequence multiple bulk samples that capture tumor diversity [6, 22]. Standard germline tools fail at low VAFs because they expect variants that appear in half or all of the reads [9]. Even specialized tools like Sniffies2 include mosaic modes that target rare alleles [23], yet they cannot distinguish somatic from germline events without normal data. Somatic discovery therefore requires dedicated tumor-normal callers that evaluate matched control samples to filter non-somatic signal [9, 24, 10].

All of the aforementioned tumor-normal callers detect SVs from reads that align to a single linear reference genome [25]. Reference bias causes alignment errors that often lead callers to misidentify germline variants as somatic SVs [26, 25, 27]. Personal assemblies [28] and pangenome references [29, 30] mitigate this bias, but they still require full read alignments that demand heavy computation [31, 25, 27]. Instead, graph-based approaches compare sequences using *k*-mers and de Bruijn graphs, which capture differences between samples directly [32, 33, 34]. Tools like novoBreak assemble reads that carry tumor-specific *k*-mers [35], while colored de Bruijn graphs represent multiple samples in a single model [34]. However, standard *k*-mers and sparse markers, such as minimizers and syncmers, have notable drawbacks [36, 37]. A single mutation disrupts up to *k* overlapping *k*-mers, while subsampled markers leave sequence gaps that miss critical variation [36, 37, 38]. Locally consistent parsing (LCP) avoids these issues by splitting sequences into variable-length *cores* that guarantee full sequence coverage [39, 40]. LCP parses shared regions into identical cores that match consistently across reads [38, 41]. A single base substitution changes at most three cores at each level [41], and iterative parsing produces longer cores that capture structural context across multiple levels [38].

Here, we introduce janus, which discovers somatic SVs in a personal pangenome built from reads generated using the PacBio HiFi platform. This pangenome is similar to a colored de Bruijn graph with LCP cores instead of *k*-mers, and includes one normal sample and one or more tumor samples from the same patient. janus first searches this graph, without any alignment, for candidate sites that are traversed only by tumor reads, and then aligns only short snippets of tumor reads around these sites to the reference genome. janus then flags germ.line variants and artifacts using a classifier that leverages features of the graph and the alignments. janus calls SVs across all tumor samples in a single run; it also leverages SV features shared across tumors to improve individual SV calls. On the CASTLE tumor-normal cell line pairs [9, 42], janus predictions showed the highest average Fl score against a consensus of 18 call sets. While achieving this accuracy, starting from FASTQ files, janus processed a pair in about 11 minutes on 16 threads, about 24 times faster than mapping the two read sets alone, while using less memory and disk space.

## 2 Methods

### Overview

janus accepts long reads from a normal sample and one or more tumor samples from the same patient (Fig. 1). After homopolymer compression [43] (collapsing runs of one base), janus partitions reads into variable-length *cores* across multiple granularity levels via locally consistent parsing (LCP) [38, 39). Core and core-pair frequencies capture sample coverage and tumor purity, defining minimum read-support thresholds. janus then constructs a colored de Bruijn-like graph [34] from these cores, tracking sample-specific read counts on each node (Fig. 1a). After standard tip clipping and bubble popping [44], janus identifies edge pairs traversed exclusively by tumor reads as candidate somatic SV sites, detecting shared sites across multiple tumors only once (Fig. 1b,c).

**Figure 1.**
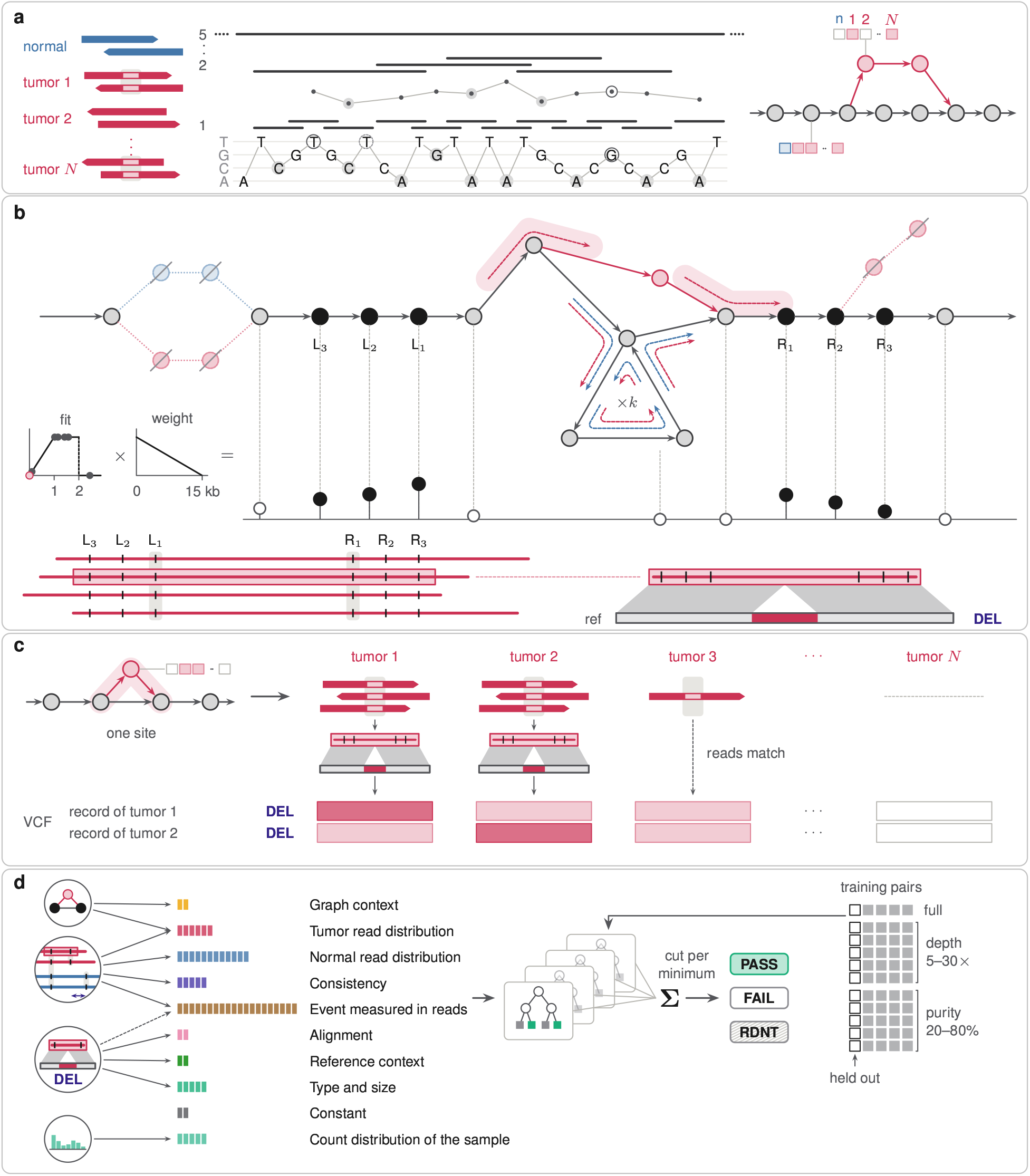
Overview of janus for one normal and *N* tumor samples. (a) The reads are parsed into LCP cores, which form a single graph with a read count per sample at each node. (b) After sequencing-error bubbles and tips are removed, a site is defined as a pair of edges traversed by tumor reads but not by normal reads; snippets of at most four tumor reads around their anchors are aligned. (c) Each site is defined once for all tumors, and each tumor with enough reads at the site gets its own VCF record; light cells: carriers found from reads spanning the site. (d) Gradient-boosted trees mark each record PASS or FAIL; duplicates are marked RDNT (redundant).

To place candidate sites on the reference genome, janus extracts read subsequences *(snippets)* bounded by normal-shared anchor cores from up to four supporting tumor reads. It maps snippets with minimap2 [43] to short reference windows seeded by shared cores, and the alignments resolve breakpoints and SV types. In a second pass, janus measures anchor-to-anchor distances across reads to detect cryptic normal support for the tumor allele. Finally, a scikit-learn [45] gradient-boosted tree classifier, trained on tumor-normal pairs at a range of depths and purities, filters candidate records using anchor distances alongside graph- and alignment-derived features (Fig. 1d).

### Core extraction

janus homopolymer-compresses each read and partitions both strands into level-5 LCP cores using LCPtools [38], indexing each core by a 64-bit sequence hash that serves as its graph node identifier.

### Graph construction

For each sample, janus counts each core and each pair of consecutive cores on the same strand of a read, skipping pairs of identical cores and cores that overlap the first or last l0bp of a read. We denote the normal sample by *So*, the tumor samples by *S*1, …, *SN*, and the count of core *x* in *S*_*S*_ by *c*_*s*_*(x)*. janus keeps a core in sample *S*_*S*_ only if *c*_*S*_*(x)* ≥ *m*_*S*,_ which removes the noise of sequencing errors, as erroneous cores occur in few reads. The minimum read support *m*_*S*_ is 2 or 3, chosen from the counts of the sample. In a tumor sample *S*_*t*_, a core with *C*_*t*_*(x)* ≥ 2 is *shared* if *c*_0_*(x)* ≥ *m*_0_ and *tumor-only* otherwise. janus sets

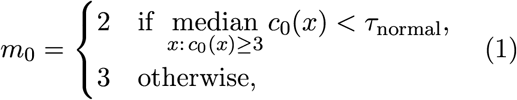

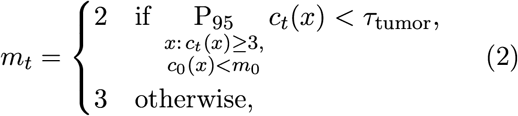

where the median grows with the depth of the normal sample, and the *95th* percentile P_95_ of the tumor-only counts is high only when the tumor sample is both deep and pure. We selected *τ*_normal_ = 20 and *τ*_tumor_ = 20 in our experiments, based on a parameter sweep over the CASTLE runs (Section Sl.l). janus then builds a graph *G*_*s*_ = *(V*_*s*_, *E*_*s*_) for each sample, with *V*_*s*_ = {x: *C*_*s*_*(x)* ≥ *m*_*s*_} and an edge (*u, v)* ∈ *E*_*s*_ for each counted pair (*u, v)* with *u, v* ∈ *V*_*s*_, and merges these graphs into the *case* graph *G* = *(V*, E) with 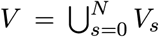 and 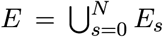 in which each node keeps its counts *c*_*S*_*(x)*. An edge is *novel* if it is not in *E*_0_ but is in some *E*_*t*_ with at least m occurrences in *S*_*t*_, where *m* = *m*_*t*_ in a pairwise run and *m* = min_*t*_ *m*_*t*_ in a joint run. On either strand, a read is the sequence of its cores that are nodes of *G*, and it traverses each edge *(u,v)* ∈ E whose endpoints are consecutive in this sequence.

From the cores that meet the minimum read support, janus computes four summary metrics to capture the depth and purity of each pair. Letting *µ*_*t*_ and *µ*_0_ denote the median counts of shared cores in the tumor and normal samples, respectively, these metrics comprise log_2_ *µ*_*t*_, log_2_ *µ*_0_, and the 5^*th*^ and 95^*th*^ percentiles of tumor-only core counts normalized by *µ*_*t*_.

### Removal of sequencing errors and small variants

A node is branching if it has multiple incoming or outgoing edges, while a chain is a maximal non-branching path *x*_1_, …, *xℓ* (*ℓ* ≥ 2). To eliminate sequencing errors, SNVs, and small indels, janus prunes tips and bubbles from the graph (Section S1.2). This pruning strategy filtered 2.2−6.2 million nodes per pair into the removed set *R* ⊂ *V* across the CASTLE pairs (Table S1).

### SV site detection

Because a somatic SV often joins pre-existing genomic cores rather than creating novel nodes in the graph, it manifests *as* a tumor-specific path through nodes shared with the normal sample. To detect these events, janus identifies pairs of edges, termed *sites*, that are traversed only by tumor reads.

Let *X* denote the endpoints of unpruned novel edges (*u, v)* where *u, v ∉. R*. We define a *locus as* either a branching node in *X* or a chain that contains a node in *X*. A read crossing a locus defines a site *σ* through the edges it enters and leaves:

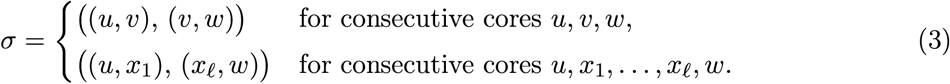

Reads that begin or end within a chain do not form a site. Letting *N*_0_ denote the set of normal read sites, janus retains a candidate site *σ* if its nodes avoid *R* and it satisfies *σ ∉. N*_0_ and max_1≤*s*≤ *N*_ *n*_*S*_*(σ)* ≥ *m*, where *n*_*S*_*(σ)* is the read count in tumor sample *s*.

For every retained site, janus records *n*_*s*_(*σ*) across all tumor samples. For each tumor sample reaching the support threshold (*n*_*s*_(*σ*) ≥ *m*), janus designates a central read r that maximizes min(*a*_*r*_, *b*_*r*_), placing the site farthest from the read ends with *a*_*r*_ and *b*_*r*_ flanking bases.

### Snippet selection

janus maps each candidate site to the reference by extracting a read snippet from the central read *r* of tumor sample *t*, bounded by flanking anchors. Let *d*(*x*) be the distance from core *x* on *r* to the site boundary. An anchor must occur uniquely on r, appear in both samples, and satisfy *c*_0_(x) ≤ 2*µ*_0_, *c*_*t*_(x) ≤ 2*µ*_*t*_, and *d*(*x*) < 15 kb. On each flank, janus selects up to three candidate anchors that maximize:

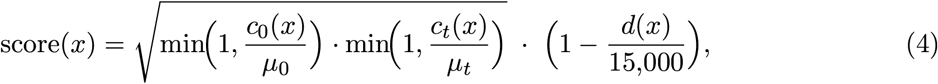

penalizing cores that are distant or poorly supported. If a flank lacks eligible cores, janus defaults to the site’s outermost core.

### Alignment of the snippets

janus aligns snippets with minimap2 [43] to short candidate reference windows rather than the whole genome (Section Sl.3). It then aligns any remaining unaligned segments (50bp to 20kb) across the whole genome to resolve distal breakpoints of translocations and non-local insertions.

### SV signatures and records

janus extracts SV signatures from split alignments and from internal indels of at least 50 bp, discarding any alignment with mapping quality below 10. Following Severus [9], each signature comprises two oriented breakpoint coordinates on the reference (Section Sl.4). Within each tumor sample, it then merges signatures of identical type and orientation with breakpoints that match within 50 bp into a single record (Section Sl.5).

### Read-level evidence

To determine whether normal reads also harbor a candidate SV, janus inspects raw reads directly using the site’s anchors. For a read *x* that carries left and right anchors *A* and *B* at positions *P*_*x*_*(A)* and *P*_*x*_*(B)*, we define the anchor distance as Δ_*x*_ = *P*_*x*_*(B)* - *P*_*x*_*(A)*. This distance matches the central read’s distance Δ_*r*_ on reads supporting the tumor allele, but shifts by approximately the SV size on alternative alleles. In a single pass across all reads, janus calculates *D*_*x*_ = Δ_*x*_ − Δ_*r*_, classifying reads with |D_*x*_| < 20 bp as supporting the tumor allele; matching normal reads signal germline variation. Because initial anchors may lie kilobases away, janus repeats this distance measurement using refined anchors closer to the event (see Section Sl.6 for their choice).

### Classification

Each record incorporates 56 features, including tumor read support, normal read *D*_*x*_ distributions, local tumor-specific node counts, and mapping quality (Section Sl.5). To account for coverage and tumor purity, janus normalizes the 11 log_2_-count features by subtracting log_2_(*µ*_*t*_ + 1) from tumor read or snippet counts and log_2_ (*µ*_0_ +1) from normal read counts. Adding *m* and the four depth/purity metrics to these features gives 61 input features per record.

In a post-processing step, janus classifies candidate records from the VCF using gradient-boosted decision trees [46, 45]. Two records of one tumor sample are duplicates if both breakpoints of one lie within 500 bp of those of the other, unless they have the same type and different orientations. In each duplicate pair, janus marks as RDNT (redundant) the record with fewer supporting tumor reads before classification and the record with the lower predicted probability after it; RDNT records are not used for training. We trained a single model across full-depth, downsampled, and 20-80% purity configurations, enabling it to generalize across variable depths and purities. janus marks a record as PASS if its predicted probability meets a support-dependent threshold (*m*) selected via held-out validation on the training pairs (Sections Sl. 7 and Sl.8).

### Joint calling of multiple tumors

In joint runs, janus initially generates and classifies records separately for each tumor using sample-specific counts, then refines calls post-classification. It clusters non-duplicate records across samples into a shared SV if the corresponding breakpoints match within 500 bp on the same chromosomes. Letting τ_*s*_ denote the probability threshold for sample S_*s*_ (Section S1.7), janus accepts a record with probability p if p ≥ τ_*s*_ + 0.2, or if p ≥ 0.1 while another tumor sample supports the same SV with p ≥ τs′. For an accepted SV, janus rescues an uncalled tumor sample as a carrier if (i) at least three other tumors report candidate records, (ii) at least one spanning read supports the event (|D_*x*_| less than 20 bp), and (iii) supporting reads comprise at least 25% of its spanning coverage. From the carrier sets of these calls, janus builds a neighbor-joining tree [47] of the tumor samples and re-decides the carriers of each SV with it: every side of every edge of the tree is a candidate carrier set, scored by the summed evidence of its members (logit p_*k*_ − logit τ_*k*_ for a tumor k with a record of the SV, − *μ* with *μ* = 0.5 otherwise) plus a prior from the number of SVs whose carriers are exactly that set; if the best candidate scores above zero, its members without an accepted record become carriers of the SV, and carriers outside it are dropped (Section S1.9).

## 3 Results

We evaluated janus using both real and simulated sequencing datasets (Section S2.1). For comparison, we ran competing PacBio HiFi-based callers using minimap2 [43] and SAMtools [48] alignments generated from identical read sets (Section S2.2) and constructed consensus truth sets using Minda [49] (Section S2.3). Sections S2.4 and S2.5 detail the simulated dataset generation and truth-set construction. To evaluate statistical significance, we computed 95% bootstrap confidence intervals [50] for the Fl score differences between janus and the best other caller across 2,000 resamples (sampling consensus SVs with replacement for recall and candidate calls for precision).

### 3.1 Accuracy analysis on real tumor-normal pairs

We first evaluated janus on five PacBio **HiFi** tumor-normal cell line pairs from the CASTLE panel [9, 42] (Table S2). The panel lacks a curated ground truth but provides an 11-call-set Minda [49] consensus of seven callers [9, 10, 11, 12, 13, 23, 24] on PacBio HiFi, ONT, and Illumina reads, requiring support from two sequencing technologies and four call sets. Because that consensus includes only four of the ten HiFi callers evaluated here, we expanded the truth set to 18 call sets by incorporating janus, our runs of all ten HiFi callers (Severus, SAVANA, nanomonsv, Sniflles2, dysgu [51], SVision-pro [52], cuteSV [53], DeBreak [54], kled [55], and SVDSS [56]; Section S2.2), and the panel’s seven ONT and Illumina sets. To ensure unbiased benchmarking, we retained the requirement of two sequencing technologies, raised the caller threshold to six, and applied uniform Minda parameters (500bp tolerance, ≥50bp length; Section S2.3). We trained the classifier on the consensus of the 17 call sets other than janus, built with the same criteria (Section Sl.7).

Using leave-one-out cross-validation over the five CASTLE pairs (Section S1.7), janus achieved the highest mean F1 score (0.887) against the 18-caller consensus, in which all callers count equally, ranking first in four pairs and trailing only the ONT call set of Severus on HCC1395 (Table S2, Fig. 2a, Table S3, Fig. S1; missed SVs analyzed in Section S3.1). While nanomonsv achieved the highest HiFi precision, janus delivered the highest recall. janus also ranked first when no caller contributed to its own truth set and when only janus was left out of the consensus (Section S3.2, Table S4). We assessed robustness by downsampling normal and tumor reads across all five CASTLE pairs to 3−5× and by generating 30× tumor purity mixtures spanning 100−20%. Scored against each pair’s full-depth consensus, janus substantially outperformed Severus, SAVANA, and nanomonsv under limited signal (Fig. 2b; Section S3.3, Table S5). Between 30× and 5× coverage, janus’s mean F1 score declined from 0.858 to 0.489, whereas Severus dropped more steeply from 0.838 to 0.265. At 20% purity, janus maintained a mean precision of 0.871 and an F1 score of 0.683, compared with 0.520 for Severus.

**Figure 2.**
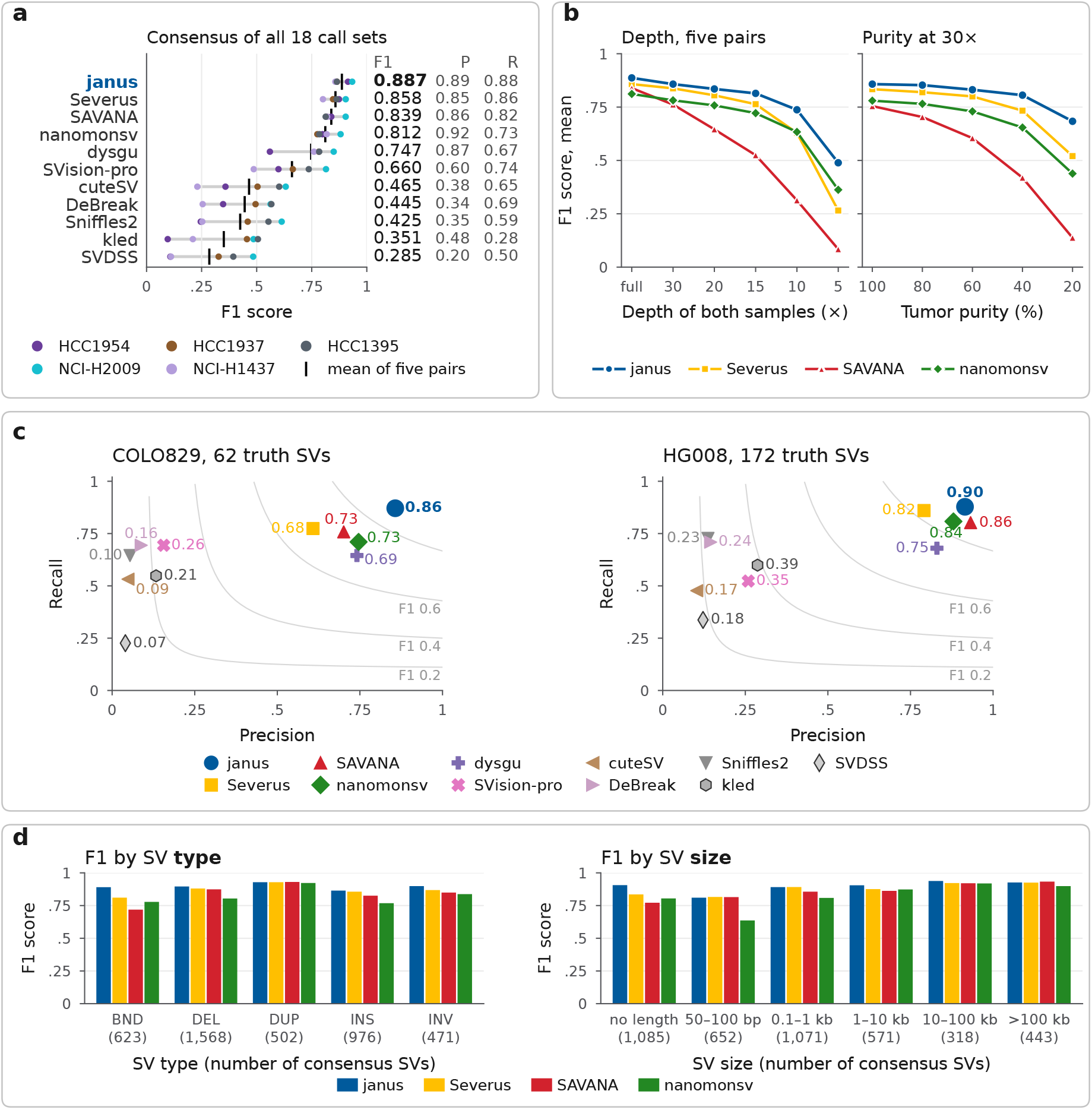
Accuracy of janus and the other callers. (a) F1 score of the 11 PacBio HiFi call sets per CASTLE pair (dots) and the mean over pairs (bar), against the consensus of all 18 call sets (P, precision; R, recall). (b) Mean F1 score at reduced depth (left) and purity (right). (c) Precision and recall on COLO829 (left) and HG008 (right), with curves of equal F1 score. (d) F1 score by SV type (left) and size (right), five pairs pooled; inversions include inverted junctions; first size class: SVs without length; consensus SVs in parentheses.

To assess generalization, we evaluated two independent benchmarks outside the training set: COLO829 (melanoma, 62 validated SVs) [57] and HG008 (pancreatic tumor, GIAB draft v0.5, 172 SVs) [58, 59]. Trained on all five CASTLE pairs, janus achieved the highest Fl score and recall among all HiFi callers on both datasets, reaching Fl scores of 0.864 on COLO829 and 0.896 on HG008 (Fig. 2c; Section S3.4, Table S6). Across pooled CASTLE pairs, janus outperformed the three published HiFi callers on breakends, deletions, insertions, inversions, and 1−100 kb SVs, whereas Severus or SAVANA led on duplications and extreme size classes (50−1,000bp and > 100kb; Fig. 2d; Table S7).

### 3.2 Calling SVs in multiple tumors within one personal pangenome

Due to the lack of multi-sample long-read tumor-normal cohorts with established ground truth, we evaluated joint calling on simulated data [60, 61, 62]: one normal genome with real germline variants (63] and 20 lineage-related tumor clones that inherit ancestral somatic SVs (Section S2.4). Across the 5,147 total simulated SVs, individual tumors harbor 1,016-1,353 variants (Fig. S2). We simulated 20× PacBio HiFi reads per genome and downsampled to l0× and 5× across five normal/tumor depth configurations (Table S8): uniform moderate depth (20/20×), uniform low depth (10/10×, 5/5×), and asymmetric designs (20/10×, 20/5×). We evaluated call sets against simulated junction coordinates using a 500bp tolerance and 50bp minimum size (Section S2.5).

We evaluated joint calling across the 20 simulated tumors in two modes: 20 separate pairwise runs against the normal sample and a single joint run spanning all 21 read sets. At 20/5× coverage, joint calling increased the median F1 score across all clones from 0.712 to 0.839 (Tables S8 and S9, Fig. 3a), driven primarily by recall (0.584 to 0.767) alongside modest precision gains (0.910 to 0.918). Joint calling maintained superior median F1 scores across all depths, though the advantage narrowed at 20/20× (0.860 versus 0.835). The gain comes largely from cross-sample classification, which rescues candidates confirmed with high confidence in another tumor (Methods, joint calling): the first pass alone reached 0.796 at 20/5×, and the clade pass added the rest, reporting 7,078 carriers of which 96% were true (Table S10). The joint run at 20/5× thus reached the median F1 score of separate runs at 20/20× (0.839 versus 0.835) with a quarter of the tumor reads. Pooling the 20 tumors into one 100× sample instead, the bulk alternative to a joint run, found 1,272 of the cohort’s 5,147 distinct SVs at 20/5×, against 2,951 for the joint run (Fig. 3c, Table S11): the pool recovered the SVs shared by six or more tumors nearly *as* well (0.78 versus 0.86) but only 4% of the 3,118 private SVs, against 45% for the joint run. Its higher precision (0.905 versus 0.685) follows from the same selectivity: a private SV has a 5% share of the pooled reads, which the classifier scores *as* an artifact, so the pool calls only the shared SVs, whereas the false calls of the joint run are mostly records of a single tumor at 5× (Table Sll). Starting from packed reads, the joint run processed all 20 tumors in 12.8min, 3.4× faster than the aggregated pairwise runs (43.7min; Fig. 3b), with only a minor memory increase over the largest pairwise run (11.9 versus 11.0GB at 20/5×; Fig. 3b). In contrast, reference alignment and sorting via minimap2 required 1 hour 58 minutes at 5/5× and an extrapolated 7.7 hours at 20/20× prior to variant calling.

**Figure 3.**
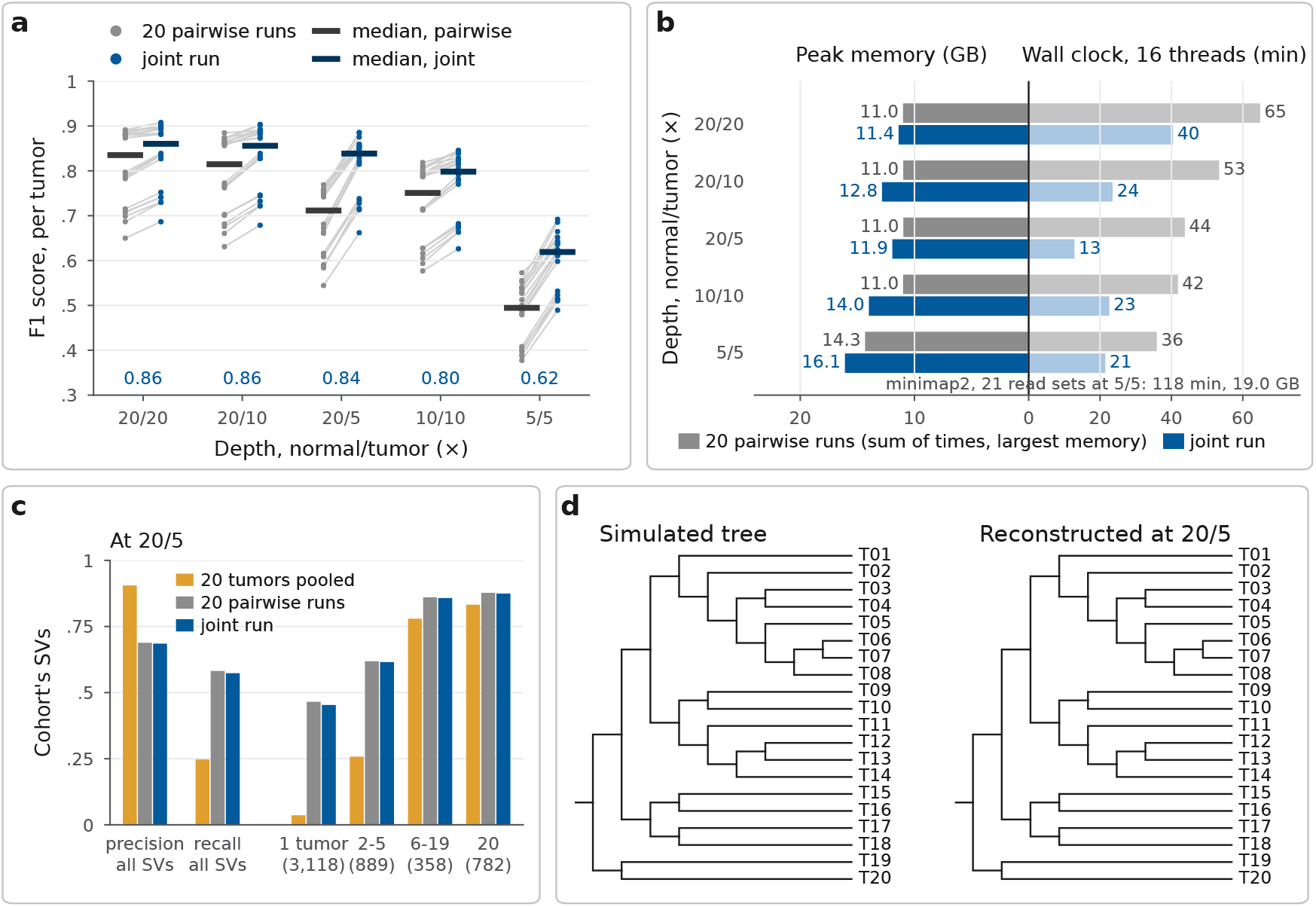
A simulated case with one normal and 20 tumors at five normal/tumor depths. (a) Fl score per tumor for the pairwise runs and the joint run with the clade pass; bars: medians over the 20 tumors; numbers: median of the joint run. (b) Wall-clock time from the packed reads on 16 threads and peak memory of the pairwise runs (sum of the times, largest memory) and of the joint run, labeled by depth. (c) Precision and recall over the cohort’s 5,147 distinct SVs at 20/5, and recall by the number of tumors carrying the SV: the 20 tumors pooled into one sample, the pairwise runs together, and the joint run. (d) The simulated tree and the tree reconstructed from the first-pass joint calls at 20/5.

The first-pass calls at 20/5× yielded 5,873 unique SVs (1,846 false positives, with 1,730 restricted to single tumors), and a neighbor-joining tree [47, 64] built on their pairwise SV differences recovered all 17 simulated splits (Robinson-Foulds distance 0 [65]; Fig. 3d), as did the trees reconstructed at the other four depth settings (Table S8).

### 3.3 Runtime and memory per pair

Mapping both read sets of HCC1954 with minimap2 and sorting the alignments took 4 hours 19 minutes with a peak memory of 19.3 GB on our workstation (Section S4.l). The complete janus run on this pair took 10 minutes 42 seconds with 10.9 GB, about 4% of this time and 56% of this memory (Table S12). With variant calling included, the alignment-based pipelines took at least 4 hours 21 minutes (cuteSV and kled). Severus and SAVANA each spent 56 minutes of their run time on calling small variants with Clair3 [66] and phasing them with LongPhase [67] (Table S13, Fig. 4a).

**Figure 4.**
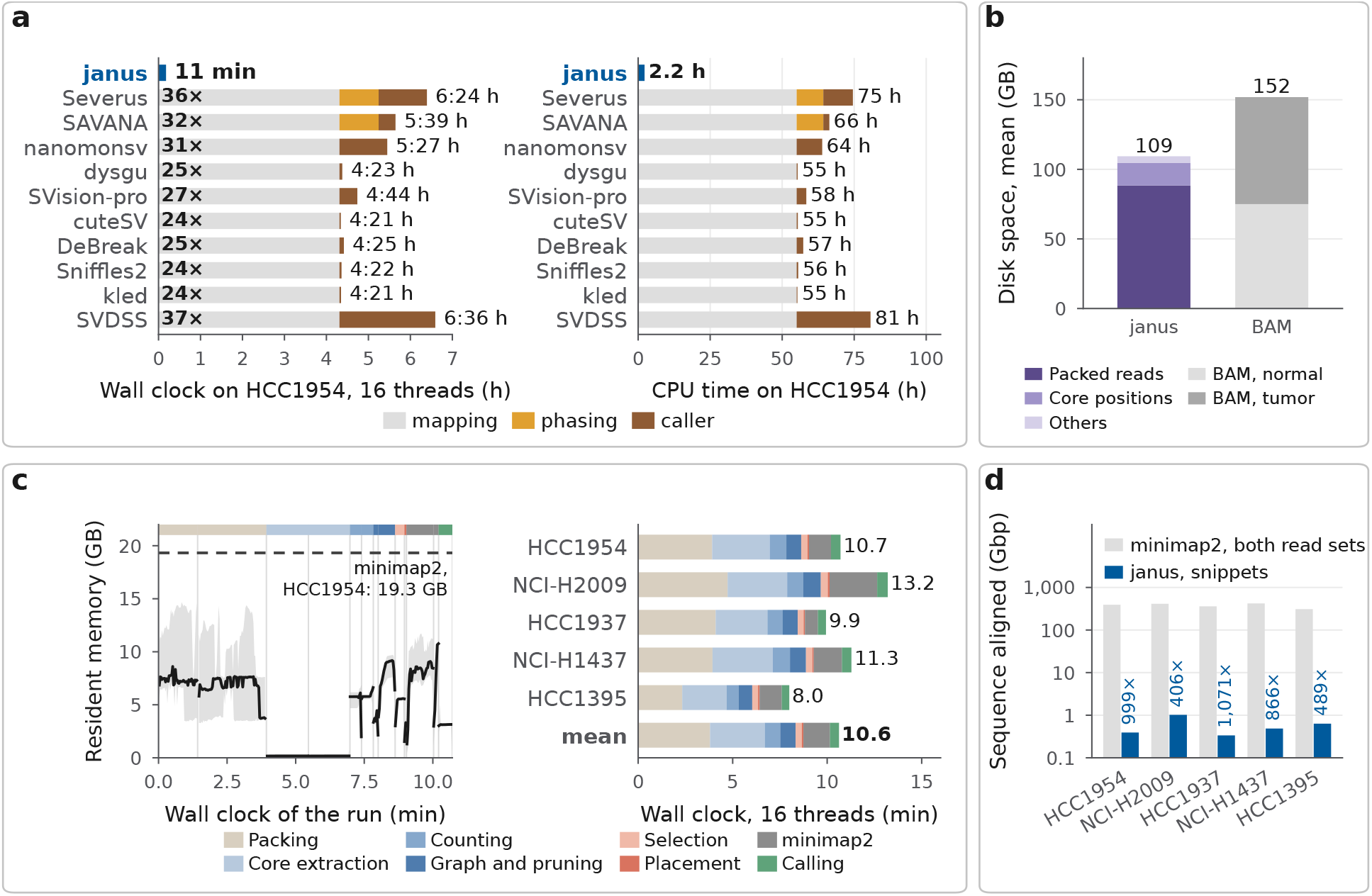
Computational cost of janus and of the alignment-based pipelines on 16 threads (SAVANA on 8 threads). (a) Wall-clock time (left) and CPU time (right) from the read sets to the calls on HCC1954, with the factor relative to janus. (b) Disk use, mean of the five pairs. (c) Resident memory over a run of janus: median (line) and range (band) of the five pairs; strip: stages, each step as wide as its median duration; dashed: minimap2 peak on HCC1954; right: wall-clock time of janus by stage, per pair and as the mean of the five pairs. (d) Sequence aligned per pair: both read sets for minimap2, snippets for janus.

Within janus, minimap2 reached a peak memory of 10.8−12.1GB when aligning the snippets or the unaligned parts of the snippets, whereas the other steps of janus used up to 9.7GB (Fig. 4c, Table S14). On disk, the intermediate files of janus require 90−121 GB per pair, most of this space for the packed reads, whereas the two sorted BAM files needed by the alignment-based callers require 117−167GB, which is on average 39% more (Fig. 4b; Section S4.2, Table S15). Starting from the FASTQ files, janus took 8 to 13 minutes per pair (Table S14, Fig. 4c). The snippets that janus aligned amounted to 0.34−1.02 Gbp per pair, which is 0.09−0.25% of the 309−420 Gbp of the two read sets of the same pair (Fig. 4d); janus did not align any read of the normal sample.

## 4 Discussion

In this work, we introduce janus, which avoids aligning all reads by building a personal pangenome, a single graph of LCP cores, built from the normal and tumor samples of a patient. Instead of searching for somatic SVs in alignments to a single reference genome, janus finds candidate sites directly in this graph and aligns only short snippets of tumor reads around them. Therefore, janus processes each tumor-normal pair in about 11 minutes, while its accuracy on the same reads is, on average, higher than that of the alignment-based callers we compared. When we reduced the sequencing depth and tumor purity of the CASTLE pairs, janus also lost less accuracy than Severus, SAVANA, and nanomonsv. The efficiency of janus matters most for large multi-sample studies, where it places all tumor samples from a patient into a single graph and processes them in a single run. For a simulated patient with 20 tumors, such a joint run was both faster and more accurate than separate janus runs for each tumor, with the largest gains at the lowest tumor depth: at 20/5×, a joint run matched separate runs at 20/20× with a quarter of the tumor reads, and it found twice as many of the cohort’s SVs as the same reads pooled into one bulk sample, which loses the SVs private to one tumor. Since a lower depth means fewer reads to sequence and to analyze, this robustness could make studies with many samples per patient cheaper.

However, janus currently builds a separate graph for each patient and uses a gradient-boosted tree classifier trained on a small cohort. A single graph of LCP cores could instead represent a whole cohort of patients, each with several tumor samples. Trained on such cohorts, deep learning models could replace the classifier and also refine the type of each SV. Reads from other long-read technologies, such as ONT, could also be added to the graph. Spatially resolved DNA sequencing [68] could locate the detected SVs and the subclones carrying them in the tissue. If pangenome-based methods scale to large cohorts, fast and accurate somatic SV calling could become a routine part of precision medicine, guiding the choice and development of targeted drugs [8].

## Acknowledgements

We thank B. ahin and K. Zambaku for suggestions during the writing of this manuscript. This work was partly supported by the European Union’s Horizon Programme for Research and Innovation under grant agreement No. 101047160 (Project BioPIM).

## LLM Usage Statement

We followed the ISCB Acceptable Use of Large Language Models Policy in this work. Claude Opus (Anthropic) and Gemini Flash (Google) were used to assist with implementing janus and to rephrase sentences and correct grammar in the manuscript. All generated code and text are inspected, reviewed, and corrected by the authors.

