## Supplementary Information for "Somatic structural variation discovery on cancer pangenome graphs"

The data shown in every figure are provided as Supplementary Data 1, one table per figure.

### S1 Implementation details of JANUS

#### S1.1 Choice of the depth thresholds

We tested $\tau_{\text{normal}} \in \{5, 10, 15, 20, 25\}$ and $\tau_{\text{tumor}} \in \{20, 30, 40, 50, 60\}$ (defined in Methods) on all CASTLE runs at full depth, reduced depth, and reduced purity (Fig. S3). Since $m_0$ and $m_t$ change only when a threshold crosses the statistic of a sample, each tested value stands for a range of thresholds with the same result on these runs: $\tau_{\text{normal}} \leq 5$, 10–13, 14–15, and 19–26; and $\tau_{\text{tumor}} = 20$, 29–32, 35–42, 43–53, and 54–60 (at $\tau_{\text{normal}} = 20$). We set $\tau_{\text{normal}} = 20$ and $\tau_{\text{tumor}} = 20$ from the ranges 19–26 and 20; across the grid, the mean F1 score of JANUS at full depth against the consensus of 18 call sets ranged from 0.884 to 0.888 (0.887 at 20/20), and 20/20 lay within 0.003 of the best combination at reduced depth and purity. The threshold affects only samples whose statistic lies near it. Depending on the sample, 56–85% of the tumor-only cores occur in only two or three reads at full depth (Fig. S4). For these statistics, JANUS assumes the worst case of the filter, $m_s = 3$, and uses only cores with $c_s(x) \geq 3$, so that the statistics do not depend on the $m_s$ it chooses.

#### S1.2 Removal of sequencing errors and small variants

JANUS identifies tips and bubbles (see Methods for definitions) on the initial graph. To find tips, JANUS follows each branch of a node with more than one outgoing edge for at most 12 nodes, as long as each node has a single incoming edge and all nodes occur in the same proper subset of the samples. If the branch ends at a node without outgoing edges, JANUS removes its nodes. JANUS does the same in the reverse direction.

To find bubbles, JANUS follows each branch of a node with at least two outgoing edges through nodes with a single outgoing edge until it reaches a node with at least two incoming edges; it follows a branch for at most five nodes, or for at most 41 nodes if its first node is missing from a sample. Two branches that end at the same node form a bubble if all nodes of one branch occur in the same proper subset of the samples and no node of the other branch occurs in exactly this subset. JANUS matches a sample-specific branch without such a counterpart with an alternative path between the same two nodes through nodes observed in all samples, which it finds by breadth-first search (depth at most the branch length plus 8, at most 256 visited nodes). JANUS extracts the sequences of both branches from reads that contain them and trims their common prefix and suffix; if the rest differs in length by less than 20 bp and has an edit distance below 20, it removes the inner nodes of the sample-specific branch as an error or a small variant (Table S1).

#### S1.3 Placement and alignment of the snippets

Given outermost anchor coordinates $p^-$ and $p^+$, and an innermost anchor span $g$, JANUS extracts the snippet $[p^- - e,\, p^+ + e]$ with padding $e = \max(1\,\text{kb},\, 3g)$, bounded by the read ends and merging any overlapping intervals. To capture distinct alleles, such as heterozygous tandem repeats, JANUS clusters supporting tumor reads by their innermost anchor distances $\delta_1 \leq \cdots \leq \delta_k$, splitting groups where $\delta_i - \delta_{i-1} > 20\,\text{bp}$. From each of the three largest clusters with at least three reads, JANUS extracts an additional snippet from its most central read.

JANUS computes the LCP cores of the reference as for the reads and indexes them by position. To define each window, JANUS requires at least three snippet cores that map uniquely to the same chromosome and strand in the reference, with diagonals of consecutive cores agreeing within 500 bp; it discards any snippet lacking an eligible window. The window spans the reference cores that define it, extended on both sides by the snippet length plus 2 kb; JANUS merges overlapping windows and aligns the snippets to them with minimap2 (v2.31; `-cx map-hifi --secondary=no`). JANUS then aligns the remaining unaligned parts of 50 bp to 20 kb to the entire reference with the short-read preset (`-cx sr --secondary=no`).

#### S1.4 SV signatures

A signature consists of two breakpoints, each a reference position with a direction; a deletion of length $L$ at position $P$, for example, is the signature $(P, +):(P + L, -)$. For split alignments, JANUS sorts the alignments of a snippet by their start on the snippet, and each consecutive pair yields a breakend if they lie on different chromosomes and an inversion if they lie on the same chromosome but on different strands. On the same chromosome and strand, the pair yields a deletion if its reference distance exceeds its distance on the snippet by at least 50 bp, an insertion if the snippet distance exceeds the reference distance by at least 50 bp and the alignments overlap by less than

50 bp on the reference, and a duplication if the second alignment starts at least 50 bp before the end of the first on the reference. Except for insertions, JANUS discards signatures with breakpoints less than 50 bp apart.

Within an alignment, JANUS takes insertions and deletions of at least 50 bp from the CIGAR string and combines those of the same type separated by fewer than 100 aligned bases. As aligners may report a tandem duplication as an insertion, an insertion of length $L \geq 500$ bp becomes a tandem duplication if the fractions $u$ and $d$ of its 15-mers found in the reference sequence of length $L$ upstream and downstream of the insertion point satisfy $0.8 \leq u + d \leq 1.25$; the duplication then starts $L\,u/(u+d)$ bases upstream of the insertion point. JANUS applies this conversion after it computes the feature values.

#### S1.5 Records, features, and output

JANUS assigns a signature to a site of its snippet whose interval between the innermost anchors lies within 200 bp of the signature; if several sites qualify, it chooses the one with the most tumor reads. JANUS discards signatures without a site, on contigs other than chromosomes 1–22, X, and Y, or at sites with fewer than $m$ tumor reads. For merging signatures into records (Methods), JANUS processes signatures in sorted order and compares each with the representative of the most recently formed record (Fig. S5). JANUS represents a record by its snippet with the most tumor reads and takes the maxima over the merged snippets as its numbers of matching and spanning normal reads.

Each record has 56 feature values, two of which are constant. They describe the tumor read support and depth at the site, normal-read evidence (the $D$ values and their dispersion), agreement of the measured distances with the event length, tumor-specific and branching nodes around the site, mapping quality and divergence of the alignment, repeat content within 150 bp of the first breakpoint, overlap of the breakpoints with annotated tandem repeats, type and size of the event, and the measurements with the second set of anchors (Fig. S6). JANUS writes all records to one VCF file, one record per row, with the feature values in the INFO field. Each sample, the normal sample included, has one column with a genotype and the number of reads of the sample with the site; the genotype is 0/1 for the tumor sample of the record and 0/0 for the other samples, whose own records are separate rows. With several tumor samples, an SV has a separate record for each tumor sample with at least $m$ reads at its site.

#### S1.6 Read-level evidence

JANUS computes $D$ (defined in Methods) for each read with an anchor on both sides of a site, using on each side the innermost of the up to three anchors present in the read, which also measures reads without the innermost anchor. The other anchors detect reads in which an anchor comes from a different locus: a read is consistent if it contains all three anchors on both sides and, on each side, the distances from the innermost anchor to the other two differ by at most 20 bp from those on the central read. The numbers of consistent tumor and normal reads, and of consistent normal reads with $|D| < 20$ bp, are features in addition to the counts over all measured reads.

JANUS also measures each signature with a second set of up to three anchors per side, which it selects on the central read before the pass: cores within 15 kb of the event and at least 20 bp outside it that occur once on the read, are present in the tumor sample, and have a normal count above zero and at most twice the median normal count. For insertions, deletions, and duplications that start within 50 bp of an annotated tandem repeat, JANUS moves these anchors to the repeat boundaries; if the total shift would exceed 10 kb, it moves each side by at most 1 kb. For duplications of at most 2 kb, it also moves them outward by the duplication length. If no anchors exist at the shifted positions, JANUS uses the unshifted ones. Each read that contains such anchors yields $D$ and the number of cores between the anchors that occur in only one of the two sequences, the read or the snippet.

The three anchors per side of the second set give up to nine anchor pairs. For each pair, JANUS compares the measured numbers of normal and of tumor reads with the largest such numbers over all pairs. Among the pairs that reach at least 80% of both maxima, the features use the pair closest to the event; if no pair qualifies, they use the pair measured in the most reads.

Among the records of the five CASTLE pairs, 56% of those that do not match the training labels have at least one normal read with $|D| < 20$ bp, compared with 1% of those that match it (Fig. S7).

#### S1.7 Training and application of the classifier

We used the `HistGradientBoostingClassifier` of scikit-learn [45] (v1.9.1; learning rate 0.05, at most 300 iterations with the default early stopping, at most 15 leaves per tree, at least 40 records per leaf, L2 regularization 1.0). The probability of a record is the mean of five models trained with different random seeds. We labeled a record true if both of its breakpoints are within 500 bp of those of an SV in the truth set, or, for insertions, if the positions are within 500 bp. The truth set for training is the Minda consensus of the 17 call sets other than JANUS (Section 3.1); the consensus of all 18 call sets was used only for evaluation. For the CASTLE benchmark, we evaluated each pair with a model trained on the other four pairs, whereas the models for HG008, COLO829, and the simulated dataset were trained on all five pairs.

The minimum read support (defined in Methods) was $m_0 = 3$ in the full-depth, 30×, and purity runs, except the purity runs of HCC1395, and $m_0 = 2$ in all other real runs. It was $m_t = 3$ in all full-depth runs, in the 30× and 100% purity runs except those of HCC1954, in the 80% purity runs of the four other CASTLE pairs, and in the 20× and 60% purity runs of NCI-H2009 and HCC1395, and $m_t = 2$ in all other real runs. In the simulated runs, $m_t$ was 3 for 19 of the 20 tumors at 20/20 and 2 otherwise.

For each value of $m$, we selected a probability threshold on held-out predictions that exclude the records the threshold applies to: a nested leave-one-out split classifies each training pair with a model trained on the other training pairs, and we chose the threshold from 0.20 to 0.80 (step 0.05) that maximizes the F1 score on these predictions. JANUS marks records at or above the threshold of their run as PASS and the others as FAIL (Fig. S8). For an insertion, the second breakpoint used for the duplicate rule (defined in Methods) is its position plus its length.

#### S1.8 Evaluation of the classifier

We scored the held-out probabilities with the area under the ROC curve (ROC-AUC) and the average precision for each CASTLE pair at every depth and purity (RDNT records excluded, training labels; Table S16). At full depth, the ROC-AUC ranged from 0.970 (HCC1954) to 0.993 (NCI-H1437) and the average precision from 0.949 to 0.970 (Fig. S9). The mean ROC-AUC stayed at 0.980 or above down to 15× and at all purities and was 0.977 at 10× and 0.962 at 5×, where the mean average precision dropped to 0.783.

We repeated the training with every choice of three CASTLE pairs (10 splits), selecting thresholds within the training pairs, and classified the two held-out pairs at all depths and purities. Each pair was among the held-out pairs in four splits, and at full depth its F1 score against the training labels varied by at most 0.049 across them: 0.874–0.882 for HCC1954, 0.914–0.929 for NCI-H2009, 0.830–0.846 for HCC1937, 0.780–0.829 for NCI-H1437, and 0.835–0.857 for HCC1395 (Fig. S10). With four training pairs, the F1 scores of HCC1954, NCI-H2009, HCC1937, and HCC1395 lay within their three-pair ranges, and that of NCI-H1437 lay above its range (0.841). At every depth and purity, the F1 score of the held-out pairs, averaged over the 10 splits, was within 0.008 of that obtained with four training pairs (Table S17).

We ran a full grid over five parameters of the gradient-boosted trees, with three values each and 243 parameter sets in total: a learning rate of 0.02, 0.05, or 0.1; at most 150, 300, or 600 iterations; at most 7, 15, or 31 leaves per tree; at least 10, 40, or 160 records per leaf; and an L2 regularization of 0, 1, or 5. We trained and evaluated every parameter set in the same way as the classifier: as the mean of five models with different random seeds, with leave-one-out cross-validation over the five CASTLE pairs, and with thresholds chosen on the nested leave-one-out split. We averaged the F1 scores over all depths and purities except 100% purity, whose runs use the same reads as the 30× runs for four of the five pairs. The best set of the grid, chosen on the held-out pairs, reached 0.780, compared with 0.777 for the chosen parameters (Table S18, Fig. S11). We also chose a parameter set for each held-out pair by nested cross-validation, using only its four training pairs: we classified each training pair with a model trained on the other three, chose the set with the highest mean F1 score over the same depths and purities, and scored this set on the held-out pair. With these sets, the mean F1 score was 0.774, compared with 0.777 for the chosen parameters (0.866 and 0.869 at full depth). A single model instead of the mean of five changed the mean F1 score at full depth from 0.869 to 0.866.

##### S1.9 Acceptance rule of the joint runs

In joint runs (described in Methods), JANUS places and aligns snippets separately for each tumor sample. For internal branching nodes, JANUS tallies reads toward $n_s(\sigma)$ only if the node also branches within the restricted subgraph $V_0 \cup V_s$, ignoring branches induced solely by other tumor samples. We classified the simulated records with the model trained on all five CASTLE pairs and compared each record with the probability threshold that corresponds to the minimum read support of its own tumor sample. We applied the same acceptance rule and carrier conditions (defined in Methods) at all five depths. Varying the margin above the threshold from 0 to 0.3 and the minimum probability of a supported record from 0.05 to 0.3 changed the median F1 score of the first pass from 0.841 to 0.858 at 20/20 and from 0.787 to 0.798 at 20/5 (Fig. S12a,b).

The clade pass (described in Methods) builds the tree from the carrier sets of the first pass as in Section S2.5, by neighbor joining on the Manhattan distances between the presence/absence vectors of the SVs over the tumor samples. The candidate carrier sets are the two sides of every edge of the unrooted tree and the set of all tumors, 75 sets for 20 tumors. A candidate $K$ scores $\sum_{k \in K} e_k + \ln \frac{n_K+1}{N+M} + \ln M$, where $e_k$ is the evidence of tumor $k$ (defined in Methods), $n_K$ the number of SVs whose first-pass carriers are exactly $K$, $N$ the number of SVs with an accepted record, and $M$ the number of candidates; the prior is zero for a set that carries an average share of the SVs, positive for a long branch, and negative for an empty one. A tumor with several records of one SV contributes its highest probability, and the carriers of an SV are attached to its accepted record with the highest probability. Varying $\mu$ from 0.3 to 2 changed the median F1 score of the clade pass from 0.854 to 0.860 at 20/20 and from 0.800 to 0.841 at 20/5 (Fig. S12c, Table S10).

#### S2 Datasets and benchmarking

##### S2.1 Datasets

We downloaded the PacBio HiFi reads of HCC1954, NCI-H2009, HCC1937, NCI-H1437, and their matched normal cell lines from the SRA (tumor: SRR28305163, SRR28305172, SRR28305185,

SRR28305179; normal: SRR28305160, SRR28305169, SRR28305182, SRR28305175). We obtained the PacBio Revio reads of HCC1395/HCC1395BL and COLO829/COLO829BL from the PacBio public datasets (`https://www.pacb.com/connect/datasets/`) and those of HG008-T and HG008-N-D from the Genome in a Bottle FTP site [58]. The published call sets of the CASTLE panel were those released with Severus [9]. The reference genome was GRCh38 with the pseudo-autosomal regions of chromosome Y masked. Fig. S13 gives the depth and the mean read length of the 14 read sets.

#### S2.2 Running the SV detection tools

We mapped the reads with minimap2 (v2.31; `-ax map-hifi -Y --secondary=no`) and sorted the alignments with SAMtools [48] (v1.21). We called small variants of the normal sample with Clair3 [66] (v2.0.3, Revio model) and phased them with LongPhase [67], which also haplotagged the tumor and normal alignment files. We ran Severus (v1.7) with default parameters, the phased VCF file, and the tandem repeat annotation. We ran SAVANA (v1.3.8) on the haplotagged alignments with `--pb`. For nanomonsv (v0.9.0), we ran `nanomonsv parse` with default parameters and `nanomonsv get` with `--qv25` and the tandem repeat annotation. We ran Sniffles2 (v2.8.0) on each sample with the tandem repeat annotation, merged the two SNF files, and kept the calls supported only by the tumor. We also ran its mosaic mode on the tumor and removed calls that start within 1 kb of a call of the same type in a normal call set made with `--minsupport 1`. This gave a mean F1 score of 0.138 on the five CASTLE pairs, compared with 0.425 in tumor–normal mode, so we did not use the mosaic mode further. We ran SVision-pro (v2.5) in its somatic mode with the HiFi preset. We ran dysgu (v1.8.8) with `--mode pacbio-revio` on both samples (`--diploid False` for the tumor), filtered the tumor calls against the normal sample with `dysgu filter`, and kept breakends and calls of at least 50 bp. For all callers, we used only PASS calls.

cuteSV (v2.1.4), DeBreak (v1.3), kled (v1.2.11), and SVDSS (v2.1.1) have no tumor–normal mode, and we ran them on each sample separately, with a minimum SV length of 50 bp where the tool offers this option. We ran cuteSV with `--max_cluster_bias_INS 1000 --diff_ratio_merging_INS 0.9 --max_cluster_bias_DEL 1000 --diff_ratio_merging_DEL 0.5`. For the normal sample we used a minimum support of 2 in cuteSV and DeBreak, a minimum cluster weight of 2 in SVDSS (4 for the tumor), and `--F2` in kled. We ran SVDSS on alignments processed by its smoothing step. We removed a tumor call if a normal call of the same type and at least 50 bp started within 500 bp (1 kb for kled) on the same chromosome. We ran Delly (v2.6.0; `delly lr -y pb`, then `delly filter -f somatic`) only on HCC1954 (F1 score 0.746); it is not one of the 18 call sets.

#### S2.3 Consensus sets and matching criteria

We built the consensus of each CASTLE pair with `minda ensemble` (`--tolerance 500 --min_size 50`) from the 10 PacBio HiFi call sets of Section S2.2, the PASS calls of JANUS, and the seven published ONT and Illumina call sets, keeping calls supported by at least two technologies and at least six call sets (Fig. S14). The other evaluations used `minda truthset` with the same parameters: we scored HG008 against the draft benchmark, COLO829 against the validated set (62 of its 68 SVs are at least 50 bp long), and the depth and purity runs against the consensus of the full-depth pair. Minda assigns the type of a consensus call by majority vote and labels most junctions that join two positions of one chromosome in inverted orientation as breakends. For the comparison by type, we counted all such junctions as inversions in the consensus and in the call sets.

#### S2.4 Generation of the simulated dataset

We derived the simulated genomes from chromosomes 1–22 and X of T2T-CHM13v2.0 with insilicoSV [60] (v0.1.9 with local modifications). The diploid normal genome carries the phased variants of HG00733 from the HGSVC3 call set on CHM13 [63], inserted into two copies of the reference, one per haplotype: 22,303 germline SVs (11,378 insertions and 10,925 deletions), 990,336 small indels, and 3,765,908 SNVs, about two thirds of them heterozygous. For the simulated data, we ran JANUS against CHM13, so the normal genome differs from the reference by the germline variants of one individual.

We generated a tree with 39 nodes using msprime [61]. Its 20 leaves are the tumor genomes, and its root, the most recent common ancestor of the tumors, descends from the normal genome. We introduced somatic variants along the branches, and each node inherits the variants of its parent. We placed each variant on a single haplotype, and the variant was heterozygous when it arose. The simulation contains 5,147 somatic SVs (1,864 deletions, 1,450 dispersed duplications, 605 tandem duplications, 581 translocations, 493 insertions, and 154 inversions), two whole-genome duplications, a cluster of complex events, 5,603 somatic small indels, and 55,320 somatic SNVs (Fig. S2). Afterward, we applied copy-number gains and losses of chromosome segments to each tumor genome, including the loss of one haplotype in some regions.

We simulated PacBio HiFi reads with Badread [62] at 20× per genome (2021 PacBio error and quality models, mean read length 17 kb), trimmed the adapters, and removed reads shorter than 100 bp; we downsampled the 10× and 5× read sets from these.

#### S2.5 Evaluation of the simulated dataset

insilicoSV reports the SVs of each genome of the simulated tree in the coordinates of its parent genome; we converted all 5,147 SVs to reference coordinates and derived the junctions each creates. We excluded SVs shorter than 50 bp, except dispersed duplications and translocations, from the truth sets and ignored calls shorter than 50 bp. A call is a true positive if both breakpoints lie within 500 bp of the two ends of a junction; precision is the fraction of calls matching a junction, and recall the fraction of SVs with at least one matched junction. In the pairwise runs at 20/20, recall was 0.72–0.76 for all SV types except translocations (0.89; Fig. S15).

For the phylogenetic analysis, we transitively merged records of different tumor samples within 500 bp into SVs. Each SV with an accepted record became a presence/absence vector over the 20 tumors, in which a tumor is present if it has an accepted record or JANUS reports it as a carrier. We built the tree from the Manhattan distances between these vectors with the neighbor-joining function of the R package ape [64]. A second reconstruction, which builds the tree directly from the most frequent compatible carrier sets, recovered 16 of the 17 splits at 20/5 (Robinson–Foulds distance 2).

For the run-time comparison, minimap2 mapped the 21 read sets one after the other, reading FASTQ streams decoded from the packed files; the time includes sorting and indexing with SAMtools.

For the pooled run, we concatenated the packed reads of the 20 tumors at 5× into one tumor sample of 98× and ran the pairwise pipeline on it with the normal sample at 20× and the same model (14.5 min on 16 threads). We scored the pooled run, the joint run, and the union of the 20 pairwise runs against the union of the 20 truth sets, after merging the accepted records within 500 bp into one call per SV; a call is true if it matches a junction of any tumor, and an SV is found if any call matches one of its junctions (Table S11).

#### S3 Accuracy on the tumor–normal pairs

On ONT data, Severus reached a mean F1 score of 0.841, whereas SAVANA and nanomonsv had lower F1 scores and precision than on HiFi (Table S3, Fig. S1a). Among the Illumina call sets, Manta and GRIPSS had higher F1 scores than the ONT call set of Sniffles2 and five HiFi call sets, while SvABA had higher F1 scores only than kled and SVDSS.

##### S3.1 Consensus SVs that were not called

Of the 4,140 SVs in the five consensus sets, 474 had no PASS call of JANUS under the matching criterion of the training labels (Section S1.7; Minda counted 456 false negatives). Of these 474 SVs, 256 had a record marked FAIL or RDNT, and for the remaining 218 JANUS wrote no record (Fig. S16).

##### S3.2 Other consensus sets

The consensus of the 17 call sets other than JANUS, built with the same criteria, contains 989, 887, 675, 215, and 1,262 SVs for HCC1954, NCI-H2009, HCC1937, NCI-H1437, and HCC1395. Against it, JANUS had a mean F1 score of 0.866, compared with 0.859 for Severus and 0.847 for SAVANA (Table S4, Fig. S1b). We also scored each of the 11 PacBio HiFi call sets against a consensus of the other 17 call sets, which for JANUS is the consensus above; these consensus sets had 988–1,044 SVs for HCC1954 and 211–221 for NCI-H1437. JANUS led all five pairs, and Severus and SAVANA reached mean F1 scores of 0.833 and 0.825 (Table S4, Fig. S1c). By SV type and size, JANUS led against the consensus without JANUS on breakends, inversions, SVs without length, and 1–10 kb SVs (Fig. S17).

##### S3.3 Reduced sequencing depth and tumor purity

To downsample a read set, we shuffled its reads and took them in this order until we reached the target number of bases, calculated for a genome size of 3.1 Gbp. A purity mixture combined tumor and normal reads in the target proportion to 30×, with a separate matched normal read set of 30×, except for HCC1395, where only 17× remained because the mixtures needed the other normal reads. We downsampled and mixed HG008 and COLO829 in the same way (Fig. S18). At 20% purity, JANUS reached F1 scores of 0.650–0.749 in the five pairs, compared with 0.473–0.562 for Severus (Table S5).

##### S3.4 COLO829, HG008, and SV types

On HG008, JANUS had the highest recall of the PacBio HiFi call sets but a lower precision than SAVANA (0.915 and 0.932; Table S6), and Table S7 gives the F1 score by SV type and size.

#### S4 Runtime and memory

##### S4.1 Test system and run times

All tools ran on a workstation with an AMD Ryzen 9 9950X CPU (16 cores, 32 hardware threads) and 196 GB of memory, using 16 threads (SAVANA on 8, as 16 threads exceeded the memory). Memory values are peak resident set sizes (RSS); we give memory and file sizes in GB.

Most of the run time of JANUS, 4.7–7.9 min per pair, went into packing the reads and extracting the cores (Table S14). Building the graph and detecting the sites took less than 2 min, and selecting, placing, and aligning the snippets took 1–3 min. Training one classifier model took 4–5 s, and the complete classification step for all 77 runs on real data (the full-depth, downsampled, and mixed read sets of the seven pairs), including the threshold selection, took 3.4 min. From 11.6–12.7 million tumor reads per pair, JANUS detected 90,000–304,000 sites, cut 26,000–75,000 snippets, and wrote 2,300–5,000 records (Fig. S19).

#### S4.2 Intermediate files

Each step of JANUS reads the files of the previous steps from disk. The packed reads and their index take most of the space, with 72–98 GB per pair, followed by the core positions on the reads with 13–18 GB, which later steps reuse instead of partitioning the reads again. The core and edge counts (one file per sample), the graph (one file for all samples, with the count of every node in every sample), and the files of the later steps (sites, snippets, reference windows, alignments) take 4–6 GB together. In total, the intermediate files of a pair are 68–77% of the size of the two sorted BAM files that the alignment-based callers require (Table S15).

### Supplementary Tables

**Supplementary Table S1:** Number of objects at each step of JANUS

| | HCC1954 | NCI-H2009 | HCC1937 | NCI-H1437 | HCC1395 |
|---|---|---|---|---|---|
| Nodes removed as tips or bubbles | 2,161,476 | 4,849,009 | 6,175,335 | 4,119,500 | 5,755,178 |
| Sites | 96,724 | 303,742 | 90,227 | 133,582 | 221,909 |
| Additional rows from distance groups | 169 | 956 | 234 | 447 | 1,482 |
| Reads from which snippets were cut | 28,260 | 74,931 | 25,847 | 33,518 | 47,650 |
| Snippets | 28,328 | 74,948 | 25,855 | 33,521 | 47,821 |
| Sequence in snippets (Mbp) | 392 | 1,018 | 335 | 486 | 631 |
| Snippets placed on the reference | 23,552 | 64,207 | 22,397 | 27,269 | 43,624 |
| Sites with an event | 3,521 | 6,685 | 2,724 | 2,918 | 6,935 |
| Records | 2,889 | 4,374 | 2,393 | 2,336 | 5,044 |
| PASS calls after classification | 1,114 | 901 | 687 | 206 | 1,210 |

Full-depth runs of the five CASTLE pairs. A snippet can contain several sites, and several snippets can carry the same event.

**Supplementary Table S2:** Accuracy of JANUS on the five CASTLE pairs

| Pair | Coverage n/t | Truth | Calls | TP | FP | FN | Precision | Recall | F1 |
|---|---|---|---|---|---|---|---|---|---|
| HCC1954 | 62.9/63.3 | 1,051 | 1,114 | 990 | 124 | 61 | 0.889 | 0.942 | 0.915 |
| NCI-H2009 | 68.1/65.1 | 896 | 901 | 839 | 62 | 57 | 0.931 | 0.936 | 0.934 |
| HCC1937 | 56.0/59.8 | 694 | 687 | 597 | 90 | 97 | 0.869 | 0.860 | 0.865 |
| NCI-H1437 | 71.1/64.5 | 221 | 206 | 183 | 23 | 38 | 0.888 | 0.828 | 0.857 |
| HCC1395 | 41.3/58.3 | 1,278 | 1,210 | 1,075 | 135 | 203 | 0.888 | 0.841 | 0.864 |

Precision, recall, and F1 score of JANUS against the consensus of 18 call sets. A model trained on the other four pairs classifies each pair. n/t, normal/tumor; TP, true positives; FP, false positives; FN, false negatives.

**Supplementary Table S3:** F1 score of the 18 call sets on the five CASTLE pairs

| Call set | Technology | HCC1954 | NCI-H2009 | HCC1937 | NCI-H1437 | HCC1395 | Precision | Recall | F1 |
|---|---|---|---|---|---|---|---|---|---|
| JANUS | PacBio HiFi | 0.915 | 0.934 | 0.865 | 0.857 | 0.864 | 0.893 | 0.882 | 0.887 |
| Severus | PacBio HiFi | 0.874 | 0.904 | 0.847 | 0.801 | 0.863 | 0.855 | 0.862 | 0.858 |
| Severus | ONT | 0.880 | 0.872 | 0.811 | 0.772 | 0.871 | 0.795 | 0.897 | 0.841 |
| SAVANA | PacBio HiFi | 0.839 | 0.904 | 0.819 | 0.820 | 0.813 | 0.863 | 0.819 | 0.839 |
| nanomonsv | PacBio HiFi | 0.799 | 0.881 | 0.776 | 0.818 | 0.785 | 0.922 | 0.728 | 0.812 |
| nanomonsv | ONT | 0.794 | 0.848 | 0.756 | 0.693 | 0.802 | 0.773 | 0.791 | 0.779 |
| dysgu | PacBio HiFi | 0.560 | 0.850 | 0.782 | 0.760 | 0.784 | 0.869 | 0.670 | 0.747 |
| SAVANA | ONT | 0.829 | 0.742 | 0.671 | 0.729 | 0.762 | 0.704 | 0.803 | 0.746 |
| SVision-pro | PacBio HiFi | 0.599 | 0.815 | 0.664 | 0.487 | 0.737 | 0.600 | 0.744 | 0.660 |
| Manta | Illumina | 0.824 | 0.302 | 0.616 | 0.546 | 0.701 | 0.546 | 0.684 | 0.598 |
| GRIPSS | Illumina | 0.810 | 0.273 | 0.509 | 0.537 | 0.682 | 0.520 | 0.647 | 0.562 |
| Sniffles2 | ONT | 0.301 | 0.637 | 0.484 | 0.344 | 0.687 | 0.438 | 0.593 | 0.490 |
| cuteSV | PacBio HiFi | 0.358 | 0.632 | 0.504 | 0.231 | 0.602 | 0.377 | 0.651 | 0.465 |
| DeBreak | PacBio HiFi | 0.348 | 0.563 | 0.495 | 0.255 | 0.567 | 0.340 | 0.693 | 0.445 |
| Sniffles2 | PacBio HiFi | 0.246 | 0.614 | 0.459 | 0.253 | 0.553 | 0.351 | 0.593 | 0.425 |
| SvABA | Illumina | 0.701 | 0.156 | 0.250 | 0.206 | 0.694 | 0.353 | 0.642 | 0.402 |
| kled | PacBio HiFi | 0.096 | 0.486 | 0.456 | 0.210 | 0.506 | 0.482 | 0.284 | 0.351 |
| SVDSS | PacBio HiFi | 0.108 | 0.484 | 0.327 | 0.111 | 0.394 | 0.205 | 0.505 | 0.285 |

F1 score per pair against the consensus of 18 call sets, and mean precision, recall, and F1 score over the five pairs, ordered by mean F1 score.

**Supplementary Table S4:** The PacBio HiFi call sets against the consensus of the 17 call sets other than JANUS and against the consensus of the other 17 call sets

| | Consensus of the 17 call sets other than JANUS | | | Consensus of the other 17 call sets | | |
|---|---|---|---|---|---|---|
| Call set | Precision | Recall | F1 | Precision | Recall | F1 |
| JANUS | 0.861 | 0.874 | 0.866 | 0.861 | 0.874 | 0.866 |
| Severus | 0.845 | 0.876 | 0.859 | 0.815 | 0.852 | 0.833 |
| SAVANA | 0.859 | 0.838 | 0.847 | 0.841 | 0.811 | 0.825 |
| nanomonsv | 0.919 | 0.745 | 0.821 | 0.905 | 0.723 | 0.802 |
| dysgu | 0.864 | 0.683 | 0.754 | 0.844 | 0.663 | 0.734 |
| SVision-pro | 0.595 | 0.757 | 0.662 | 0.580 | 0.732 | 0.643 |
| cuteSV | 0.373 | 0.662 | 0.466 | 0.364 | 0.639 | 0.452 |
| DeBreak | 0.339 | 0.707 | 0.448 | 0.331 | 0.685 | 0.436 |
| Sniffles2 | 0.350 | 0.605 | 0.428 | 0.341 | 0.585 | 0.415 |
| kled | 0.481 | 0.290 | 0.355 | 0.469 | 0.280 | 0.344 |
| SVDSS | 0.203 | 0.509 | 0.284 | 0.198 | 0.496 | 0.277 |

Mean precision, recall, and F1 score over the five CASTLE pairs.

**Supplementary Table S5:** Accuracy at reduced sequencing depth and tumor purity

| Depth | JANUS | | | | Severus | SAVANA | nanomonsv | JANUS − best other |
|---|---|---|---|---|---|---|---|---|
| | Precision | Recall | F1 | 95% interval | F1 | F1 | F1 | F1 [95% interval] |
| Full | 0.893 | 0.882 | 0.887 | 0.878–0.897 | 0.858 | 0.839 | 0.812 | +0.030 [+0.018, +0.041] vs. Severus |
| 30× | 0.850 | 0.867 | 0.858 | 0.848–0.867 | 0.838 | 0.762 | 0.782 | +0.020 [+0.006, +0.033] vs. Severus |
| 20× | 0.841 | 0.838 | 0.835 | 0.824–0.845 | 0.805 | 0.646 | 0.758 | +0.030 [+0.017, +0.042] vs. Severus |
| 15× | 0.846 | 0.795 | 0.814 | 0.803–0.825 | 0.764 | 0.525 | 0.722 | +0.051 [+0.037, +0.064] vs. Severus |
| 10× | 0.786 | 0.704 | 0.737 | 0.725–0.749 | 0.629 | 0.313 | 0.634 | +0.104 [+0.089, +0.119] vs. nanomonsv |
| 5× | 0.683 | 0.395 | 0.489 | 0.473–0.504 | 0.265 | 0.084 | 0.362 | +0.127 [+0.110, +0.143] vs. nanomonsv |
| Purity (30×) | | | | | | | | |
| 100% | 0.850 | 0.867 | 0.858 | 0.848–0.867 | 0.834 | 0.755 | 0.780 | +0.024 [+0.011, +0.036] vs. Severus |
| 80% | 0.858 | 0.850 | 0.853 | 0.843–0.862 | 0.820 | 0.703 | 0.765 | +0.033 [+0.021, +0.046] vs. Severus |
| 60% | 0.847 | 0.821 | 0.831 | 0.821–0.842 | 0.800 | 0.605 | 0.730 | +0.031 [+0.019, +0.045] vs. Severus |
| 40% | 0.855 | 0.765 | 0.806 | 0.795–0.816 | 0.733 | 0.419 | 0.654 | +0.073 [+0.060, +0.086] vs. Severus |
| 20% | 0.871 | 0.565 | 0.683 | 0.670–0.697 | 0.520 | 0.138 | 0.438 | +0.163 [+0.147, +0.178] vs. Severus |

Mean over the five CASTLE pairs against the consensus of 18 call sets of the full-depth pair. 95% interval: bootstrap interval of the mean F1 score of JANUS; JANUS − best other: difference from the caller with the highest mean F1 score, with its bootstrap 95% interval (2,000 resamples).

**Supplementary Table S6:** Accuracy on COLO829 and HG008

| | COLO829 | | | | HG008 | | | |
|---|---|---|---|---|---|---|---|---|
| Call set | Calls | Precision | Recall | F1 | Calls | Precision | Recall | F1 |
| JANUS | 63 | 0.857 | 0.871 | **0.864** | 165 | 0.915 | 0.878 | **0.896** |
| SAVANA | 67 | 0.701 | 0.758 | 0.729 | 148 | 0.932 | 0.802 | 0.863 |
| nanomonsv | 59 | 0.746 | 0.710 | 0.727 | 158 | 0.880 | 0.808 | 0.842 |
| dysgu | 54 | 0.741 | 0.645 | 0.690 | 141 | 0.830 | 0.680 | 0.748 |
| Severus | 79 | 0.608 | 0.774 | 0.681 | 187 | 0.791 | 0.860 | 0.825 |
| SVision-pro | 275 | 0.156 | 0.694 | 0.255 | 348 | 0.259 | 0.523 | 0.346 |
| kled | 256 | 0.133 | 0.548 | 0.214 | 359 | 0.287 | 0.599 | 0.388 |
| DeBreak | 491 | 0.088 | 0.694 | 0.156 | 843 | 0.145 | 0.709 | 0.240 |
| Sniffles2 | 743 | 0.054 | 0.645 | 0.099 | 912 | 0.137 | 0.727 | 0.231 |
| cuteSV | 689 | 0.048 | 0.532 | 0.088 | 794 | 0.103 | 0.477 | 0.170 |
| SVDSS | 352 | 0.040 | 0.226 | 0.068 | 476 | 0.122 | 0.337 | 0.179 |
| JANUS − best other, F1 | +0.135 [+0.035, +0.237] vs. SAVANA | | | | +0.034 [-0.023, +0.087] vs. SAVANA | | | |

COLO829 (62 validated somatic SVs of at least 50 bp) and HG008 (draft benchmark v0.5), scored with Minda; neither pair was part of the training data. Rows ordered by F1 score on COLO829; bold: highest F1 score. Last row: difference from the call set with the next highest F1 score, with its bootstrap 95% interval.

**Supplementary Table S7:** F1 score by SV type and size on the five CASTLE pairs

| | Class | Consensus SVs | JANUS | Severus | SAVANA | nanomonsv |
|---|---|---|---|---|---|---|
| Type | Breakend | 623 | **0.891** | 0.811 | 0.719 | 0.778 |
| | Deletion | 1,568 | **0.896** | 0.880 | 0.874 | 0.804 |
| | Duplication | 502 | 0.929 | 0.929 | **0.930** | 0.923 |
| | Insertion | 976 | **0.864** | 0.856 | 0.826 | 0.768 |
| | Inversion | 471 | **0.899** | 0.868 | 0.849 | 0.838 |
| Size | no length | 1,085 | **0.906** | 0.835 | 0.772 | 0.804 |
| | 50–100 bp | 652 | 0.810 | **0.816** | 0.815 | 0.636 |
| | 0.1–1 kb | 1,071 | 0.891 | **0.892** | 0.856 | 0.809 |
| | 1–10 kb | 571 | **0.905** | 0.876 | 0.862 | 0.873 |
| | 10–100 kb | 318 | **0.938** | 0.922 | 0.921 | 0.919 |
| | >100 kb | 443 | 0.927 | 0.925 | **0.933** | 0.899 |

Five pairs pooled, consensus of 18 call sets; bold: highest of each row.

**Supplementary Table S8:** Pairwise and joint runs on the simulated case

| | Pairwise runs | | | Joint run, first pass | | | Joint run, clade pass | | | Tree | | Wall clock (min) | | Peak RSS (GB) | |
|---|---|---|---|---|---|---|---|---|---|---|---|---|---|---|---|
| Depth n/t | Precision | Recall | F1 | Precision | Recall | F1 | Precision | Recall | F1 | Splits | RF | pairwise | joint | pairwise | joint |
| 20/20 | 0.891 | 0.786 | 0.835 | 0.907 | 0.803 | 0.852 | 0.896 | 0.829 | 0.860 | 17/17 | 0 | 64.9 | 40.4 | 11.0 | 11.4 |
| 20/10 | 0.894 | 0.747 | 0.815 | 0.904 | 0.781 | 0.839 | 0.896 | 0.819 | 0.856 | 17/17 | 0 | 53.4 | 23.5 | 11.0 | 12.8 |
| 20/5 | 0.910 | 0.584 | 0.712 | 0.922 | 0.697 | 0.796 | 0.918 | 0.767 | 0.839 | 17/17 | 0 | 43.7 | 12.8 | 11.0 | 11.9 |
| 10/10 | 0.824 | 0.689 | 0.751 | 0.825 | 0.743 | 0.785 | 0.819 | 0.772 | 0.798 | 17/17 | 0 | 41.8 | 22.6 | 11.0 | 14.0 |
| 5/5 | 0.500 | 0.486 | 0.495 | 0.587 | 0.611 | 0.601 | 0.602 | 0.628 | 0.619 | 17/17 | 0 | 35.9 | 21.5 | 14.3 | 16.1 |

Precision, recall, and F1 score: medians over the 20 tumors. First pass: records and read-supported carriers; clade pass: carriers re-decided with the tree (described in Methods). Splits: splits of the simulated tree found in the tree reconstructed from the first-pass calls; RF: Robinson–Foulds distance. Wall-clock time and peak memory on 16 threads from the packed reads (pairwise: sum of the times, largest memory). minimap2 mapped the 21 read sets at 5/5 in 117.6 min with 19.0 GB.

**Supplementary Table S9:** F1 score of each simulated tumor in the pairwise runs and the two passes of the joint run

| | 20/20 | | | 20/10 | | | 20/5 | | | 10/10 | | | 5/5 | | |
|---|---|---|---|---|---|---|---|---|---|---|---|---|---|---|---|
| Tumor | pairwise | first | clade | pairwise | first | clade | pairwise | first | clade | pairwise | first | clade | pairwise | first | clade |
| T01 | 0.879 | 0.889 | 0.895 | 0.866 | 0.876 | 0.886 | 0.747 | 0.825 | 0.852 | 0.796 | 0.817 | 0.826 | 0.540 | 0.618 | 0.639 |
| T02 | 0.882 | 0.895 | 0.903 | 0.865 | 0.877 | 0.893 | 0.756 | 0.848 | 0.885 | 0.794 | 0.826 | 0.839 | 0.535 | 0.644 | 0.665 |
| T03 | 0.891 | 0.904 | 0.908 | 0.871 | 0.893 | 0.902 | 0.753 | 0.842 | 0.875 | 0.809 | 0.841 | 0.845 | 0.540 | 0.662 | 0.685 |
| T04 | 0.884 | 0.897 | 0.902 | 0.872 | 0.892 | 0.904 | 0.759 | 0.849 | 0.885 | 0.803 | 0.838 | 0.845 | 0.556 | 0.665 | 0.692 |
| T05 | 0.797 | 0.822 | 0.839 | 0.766 | 0.803 | 0.833 | 0.672 | 0.759 | 0.823 | 0.715 | 0.765 | 0.791 | 0.481 | 0.589 | 0.612 |
| T06 | 0.786 | 0.811 | 0.834 | 0.762 | 0.794 | 0.828 | 0.661 | 0.751 | 0.819 | 0.714 | 0.751 | 0.779 | 0.479 | 0.582 | 0.598 |
| T07 | 0.783 | 0.801 | 0.827 | 0.767 | 0.792 | 0.828 | 0.664 | 0.762 | 0.815 | 0.712 | 0.749 | 0.770 | 0.481 | 0.600 | 0.611 |
| T08 | 0.791 | 0.810 | 0.837 | 0.773 | 0.801 | 0.839 | 0.681 | 0.770 | 0.830 | 0.713 | 0.752 | 0.781 | 0.489 | 0.608 | 0.624 |
| T09 | 0.650 | 0.670 | 0.687 | 0.631 | 0.656 | 0.679 | 0.544 | 0.623 | 0.662 | 0.577 | 0.607 | 0.626 | 0.377 | 0.481 | 0.489 |
| T10 | 0.699 | 0.717 | 0.730 | 0.676 | 0.703 | 0.722 | 0.590 | 0.672 | 0.714 | 0.605 | 0.649 | 0.663 | 0.387 | 0.499 | 0.516 |
| T11 | 0.708 | 0.730 | 0.741 | 0.703 | 0.726 | 0.744 | 0.609 | 0.694 | 0.734 | 0.627 | 0.665 | 0.677 | 0.403 | 0.499 | 0.513 |
| T12 | 0.687 | 0.711 | 0.730 | 0.661 | 0.695 | 0.722 | 0.588 | 0.668 | 0.716 | 0.593 | 0.643 | 0.664 | 0.390 | 0.499 | 0.510 |
| T13 | 0.715 | 0.739 | 0.752 | 0.701 | 0.735 | 0.746 | 0.616 | 0.701 | 0.738 | 0.628 | 0.675 | 0.682 | 0.408 | 0.527 | 0.532 |
| T14 | 0.707 | 0.728 | 0.741 | 0.682 | 0.709 | 0.732 | 0.584 | 0.683 | 0.728 | 0.615 | 0.657 | 0.673 | 0.398 | 0.509 | 0.524 |
| T15 | 0.873 | 0.883 | 0.882 | 0.860 | 0.879 | 0.882 | 0.759 | 0.829 | 0.859 | 0.790 | 0.820 | 0.823 | 0.527 | 0.620 | 0.637 |
| T16 | 0.878 | 0.891 | 0.893 | 0.856 | 0.877 | 0.885 | 0.754 | 0.829 | 0.859 | 0.787 | 0.814 | 0.818 | 0.529 | 0.624 | 0.636 |
| T17 | 0.887 | 0.900 | 0.897 | 0.885 | 0.890 | 0.890 | 0.769 | 0.832 | 0.853 | 0.818 | 0.836 | 0.839 | 0.573 | 0.633 | 0.644 |
| T18 | 0.890 | 0.899 | 0.896 | 0.874 | 0.885 | 0.888 | 0.758 | 0.833 | 0.854 | 0.811 | 0.827 | 0.829 | 0.552 | 0.639 | 0.652 |
| T19 | 0.880 | 0.889 | 0.882 | 0.860 | 0.875 | 0.875 | 0.742 | 0.822 | 0.847 | 0.787 | 0.813 | 0.814 | 0.513 | 0.598 | 0.621 |
| T20 | 0.880 | 0.892 | 0.884 | 0.863 | 0.875 | 0.873 | 0.745 | 0.827 | 0.854 | 0.791 | 0.805 | 0.805 | 0.500 | 0.603 | 0.617 |

Column groups give the normal/tumor depth; pairwise: separate runs; first, clade: first pass and clade pass of the joint run.

**Supplementary Table S10:** Carriers reported by the first pass and the clade pass of the joint run

| | Carriers, first pass | | | Carriers, clade pass | | | Median F1 of the clade pass by $\mu$ | | | | |
|---|---|---|---|---|---|---|---|---|---|---|---|
| Depth n/t | Reported | True | Precision | Reported | True | Precision | 0.3 | 0.5 | 1.0 | 1.5 | 2.0 |
| 20/20 | 1,097 | 988 | 0.901 | 1,798 | 1,492 | 0.830 | 0.854 | 0.860 | 0.857 | 0.856 | 0.855 |
| 20/10 | 2,103 | 1,951 | 0.928 | 3,066 | 2,790 | 0.910 | 0.855 | 0.856 | 0.849 | 0.848 | 0.850 |
| 20/5 | 4,800 | 4,652 | 0.969 | 7,078 | 6,794 | 0.960 | 0.841 | 0.839 | 0.829 | 0.808 | 0.800 |
| 10/10 | 1,979 | 1,803 | 0.911 | 2,570 | 2,266 | 0.882 | 0.799 | 0.798 | 0.793 | 0.791 | 0.792 |
| 5/5 | 5,448 | 4,014 | 0.737 | 5,206 | 4,317 | 0.829 | 0.625 | 0.619 | 0.605 | 0.583 | 0.575 |

Carriers: tumor samples without an accepted record that are reported for an SV, summed over the 20 tumors; True: carriers that match a simulated SV of that tumor. Right: median F1 score over the 20 tumors of the clade pass for each value of the penalty $\mu$; the paper uses 0.5.

**Supplementary Table S11:** The 20 tumors pooled into one sample, the pairwise runs, and the joint run against the SVs of the whole cohort at 20/5

| | | | | | | | Recall by the number of tumors carrying the SV | | | |
|---|---|---|---|---|---|---|---|---|---|---|
| Run | Calls | True | SVs found | Precision | Recall | F1 | 1 (3,118) | 2-5 (889) | 6-19 (358) | 20 (782) |
| 20 tumors pooled into one sample | 1,858 | 1,682 | 1,272 | 0.905 | 0.247 | 0.388 | 0.036 | 0.258 | 0.779 | 0.832 |
| 20 pairwise runs | 5,978 | 4,115 | 2,994 | 0.688 | 0.582 | 0.631 | 0.465 | 0.619 | 0.860 | 0.877 |
| Joint run | 5,874 | 4,025 | 2,951 | 0.685 | 0.573 | 0.624 | 0.453 | 0.615 | 0.858 | 0.875 |

Accepted records merged within 500 bp into one call per SV and scored against the union of the 20 truth sets (5,147 distinct SVs; Section S2.5). SVs found: distinct SVs with a matching call. Right: recall of the SVs carried by the given number of tumors (number of such SVs in parentheses).

**Supplementary Table S12:** Mapping time of the two read sets of each pair

| Pair | Reads (Gbp) | Snippets (Gbp) | Ratio | Mapping (h:mm) | Ratio |
|---|---|---|---|---|---|
| HCC1954 | 391 | 0.39 | 999 | 4:19 | 24.2 |
| NCI-H2009 | 413 | 1.02 | 406 | 4:44† | 21.5† |
| HCC1937 | 359 | 0.34 | 1,071 | 4:07† | 24.8† |
| NCI-H1437 | 420 | 0.49 | 866 | 4:49† | 25.6† |
| HCC1395 | 309 | 0.63 | 489 | 3:32† | 26.5† |

minimap2 on 16 threads, including sorting. †Estimated from the amount of sequence at the mean throughput of HCC1954 and COLO829 (39.6 and 42.8 s per Gbp). Snippets: sequence of the snippets aligned by JANUS, and the ratio of the reads to it; the second ratio is the mapping time divided by the run time of JANUS (Table S14).

**Supplementary Table S13:** Run time and memory of the pipelines on HCC1954

| Call set | Mapping | Phasing | Calling | Total | Factor | CPU (h) | Peak RSS (GB) |
|---|---|---|---|---|---|---|---|
| JANUS | – | – | – | 0:11 | 1 | 2.2 | 10.9 |
| cuteSV | 4:19 | – | 0:02 | 4:21 | 24 | 55 | 57.2 |
| kled | 4:19 | – | 0:02 | 4:21 | 24 | 55 | 40.7 |
| Sniffles2 | 4:19 | – | 0:03 | 4:22 | 24 | 56 | 19.3 |
| dysgu | 4:19 | – | 0:04 | 4:23 | 25 | 55 | 54.0 |
| DeBreak | 4:19 | – | 0:06 | 4:25 | 25 | 57 | 19.3 |
| SVision-pro | 4:19 | – | 0:26 | 4:44 | 27 | 58 | 45.5 |
| nanomonsv | 4:19 | – | 1:08 | 5:27 | 31 | 64 | 19.3 |
| SAVANA | 4:19 | 0:56 | 0:24 | 5:39 | 32 | 66 | 62.9 |
| Severus | 4:19 | 0:56 | 1:09 | 6:24 | 36 | 75 | 53.3 |
| SVDSS | 4:19 | – | 2:17 | 6:36 | 37 | 81 | 99.6 |

Wall clock (h:mm) from the read sets to the calls on 16 threads (SAVANA on 8). Factor: total time relative to JANUS. Peak RSS: largest over mapping, phasing, and calling.

**Supplementary Table S14:** Run time and peak memory of JANUS per stage

| Stage | HCC1954 | NCI-H2009 | HCC1937 | NCI-H1437 | HCC1395 | Peak RSS (GB) |
|---|---|---|---|---|---|---|
| Packing | 3:55 | 4:44 | 4:06 | 3:56 | 2:20 | 4.3–5.5 |
| Core extraction | 3:03 | 3:09 | 2:44 | 3:12 | 2:21 | 0.2 |
| Counting | 0:53 | 0:51 | 0:49 | 0:54 | 0:38 | 6.6–7.4 |
| Graph and pruning | 0:47 | 0:55 | 0:48 | 0:51 | 0:43 | 7.6–9.7 |
| Selection | 0:20 | 0:22 | 0:19 | 0:20 | 0:19 | 5.5–5.7 |
| Placement | 0:04 | 0:06 | 0:04 | 0:04 | 0:05 | 4.2–4.9 |
| minimap2 | 1:10 | 2:31 | 0:41 | 1:29 | 1:10 | 10.8–12.1 |
| Calling | 0:30 | 0:33 | 0:26 | 0:30 | 0:24 | 5.0–5.1 |
| Total | 10:42 | 13:12 | 9:56 | 11:17 | 8:00 | 10.8–12.1 |

Wall-clock time (m:ss) per stage on 16 threads and range of peak resident memory over the five pairs. RSS, resident set size.

**Supplementary Table S15:** Size of the intermediate files of JANUS

| Pair | Packed reads | Core positions | Counts | Graph | Snippets, alignments, other | Total | BAM files |
|---|---|---|---|---|---|---|---|
| HCC1954 | 91.4 | 16.7 | 2.2 | 1.0 | 1.1 | 112.5 | 165 |
| NCI-H2009 | 96.5 | 17.6 | 2.1 | 1.0 | 2.8 | 120.1 | 167 |
| HCC1937 | 83.9 | 15.3 | 2.1 | 1.0 | 1.1 | 103.3 | 144 |
| NCI-H1437 | 98.1 | 17.9 | 2.1 | 1.0 | 1.4 | 120.7 | 167 |
| HCC1395 | 72.2 | 13.2 | 2.1 | 1.0 | 1.9 | 90.3 | 117 |

Intermediate files of JANUS per pair and the two sorted BAM files that an alignment-based caller needs, in GB ($2^{30}$ bytes). Packed reads include their index.

**Supplementary Table S16:** Threshold-free evaluation of the classifier

| Setting | full | 30× | 20× | 15× | 10× | 5× |
|---|---|---|---|---|---|---|
| ROC-AUC | 0.981 | 0.980 | 0.980 | 0.980 | 0.977 | 0.962 |
| Average precision | 0.958 | 0.948 | 0.938 | 0.931 | 0.909 | 0.783 |
| Purity (30×) | 100% | 80% | 60% | 40% | 20% | |
| ROC-AUC | 0.980 | 0.980 | 0.982 | 0.984 | 0.987 | |
| Average precision | 0.947 | 0.948 | 0.947 | 0.942 | 0.939 | |

Mean over the five CASTLE pairs of the held-out probabilities, each pair classified by the model trained on the other four (RDNT records excluded, training labels). ROC-AUC, area under the receiver operating characteristic curve.

**Supplementary Table S17:** Training on three instead of four CASTLE pairs

| Setting | full | 30× | 20× | 15× | 10× | 5× |
|---|---|---|---|---|---|---|
| Four training pairs | 0.869 | 0.854 | 0.833 | 0.814 | 0.738 | 0.492 |
| Three training pairs, mean | 0.861 | 0.850 | 0.828 | 0.809 | 0.736 | 0.492 |
| Three training pairs, range | 0.812–0.905 | 0.807–0.897 | 0.767–0.884 | 0.760–0.860 | 0.665–0.792 | 0.440–0.545 |
| Purity (30×) | 100% | 80% | 60% | 40% | 20% | |
| Four training pairs | 0.854 | 0.849 | 0.829 | 0.805 | 0.686 | |
| Three training pairs, mean | 0.849 | 0.844 | 0.825 | 0.800 | 0.682 | |
| Three training pairs, range | 0.807–0.897 | 0.800–0.894 | 0.770–0.883 | 0.759–0.844 | 0.651–0.721 | |

Mean F1 score of the held-out CASTLE pairs (training labels) with four training pairs (leave-one-out) and with three (all 10 splits). The range is that of the mean of the two held-out pairs over the 10 splits.

**Supplementary Table S18:** Full parameter grid of the gradient-boosted trees

| Parameter set | Full | 20× | 5× | 20% purity | All |
|---|---|---|---|---|---|
| Parameters used in this paper | 0.869 | 0.833 | 0.492 | 0.686 | 0.777 |
| scikit-learn, 243 sets | | | | | |
| Best set (chosen on the held-out pairs) | 0.867 | 0.836 | 0.511 | 0.688 | 0.780 |
| Sets chosen by nested cross-validation | 0.866 | 0.827 | 0.486 | 0.684 | 0.774 |
| Range of the 243 sets | 0.832–0.873 | 0.819–0.836 | 0.424–0.511 | 0.678–0.688 | 0.763–0.780 |

Mean F1 score of the five CASTLE pairs (training labels) at full depth, at 20×, at 5×, at 20% purity, and over all depths and purities except 100% purity (All). Nested cross-validation chooses a set for each held-out pair on its four training pairs and can therefore combine up to five sets.

### Supplementary Figures

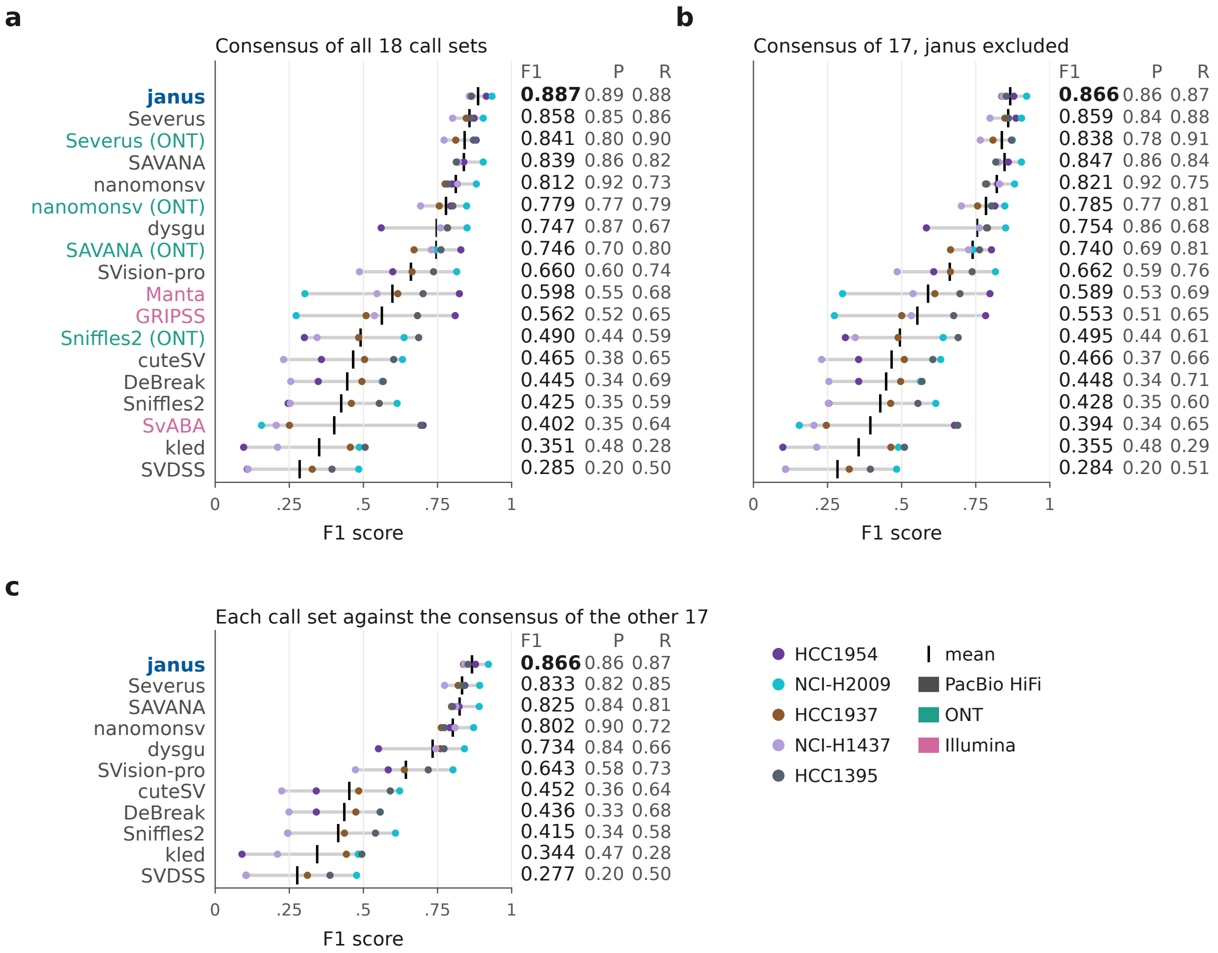

**Supplementary Figure S1:** F1 score of the call sets in each CASTLE pair (dots) and its mean (bar); columns: mean F1 score, precision (P), recall (R). (a) All 18 call sets against the consensus of 18 call sets and (b) of the 17 call sets other than JANUS; name colors give the sequencing technology. (c) Each PacBio HiFi call set against the consensus of the other 17 call sets.

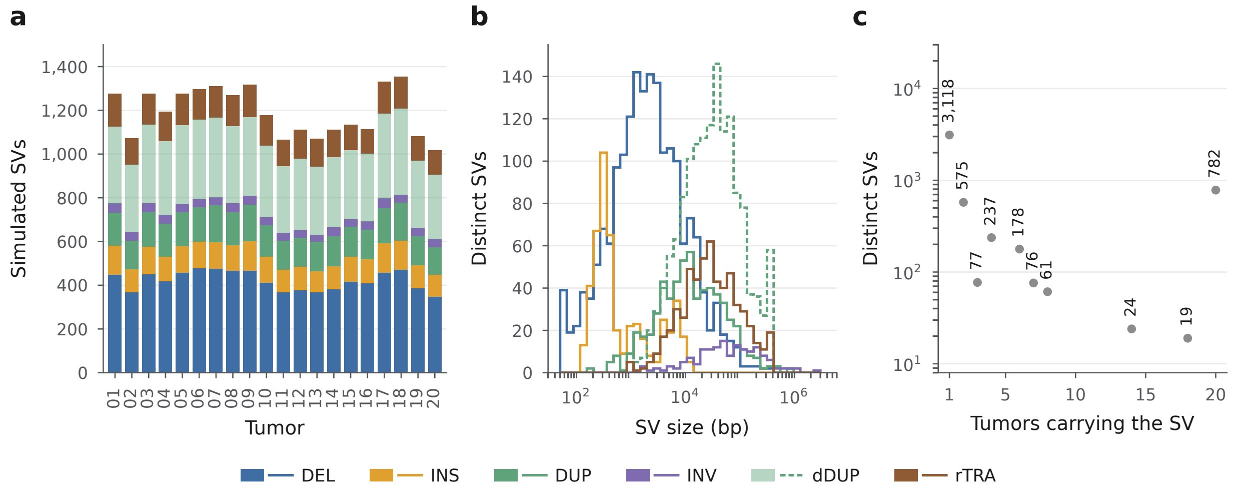

**Supplementary Figure S2:** The simulated dataset. (a) SVs of each tumor by type (DUP, tandem duplication; dDUP, dispersed duplication; rTRA, translocation). (b) Size distribution of the 5,147 simulated SVs by type. (c) Number of SVs by the number of tumors that carry them.

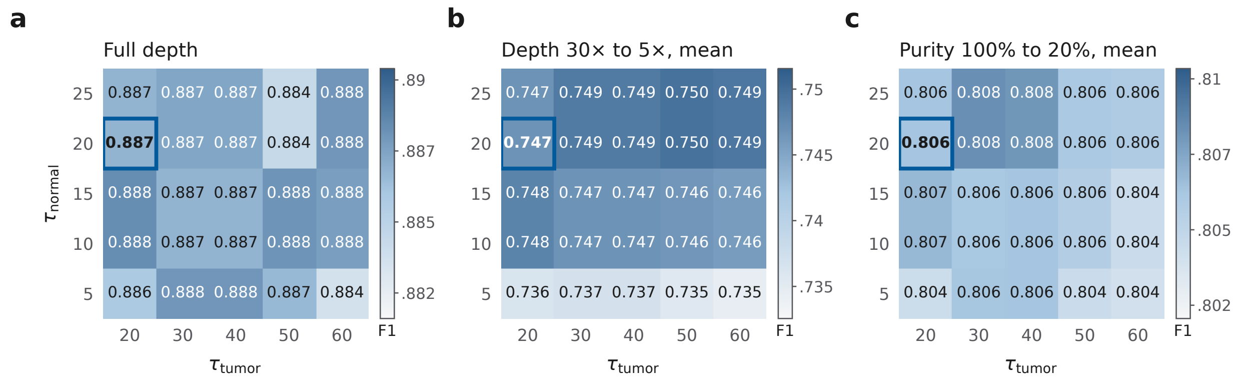

**Supplementary Figure S3:** Mean F1 score of JANUS on the five CASTLE pairs for each combination of depth thresholds, against the consensus of 18 call sets of the full-depth pair: at full depth (a), averaged over the reduced depths from 30× to 5× (b), and averaged over the tumor purities from 100% to 20% (c). The chosen thresholds, $\tau_{normal} = \tau_{tumor} = 20$, are outlined; each panel has its own color scale.

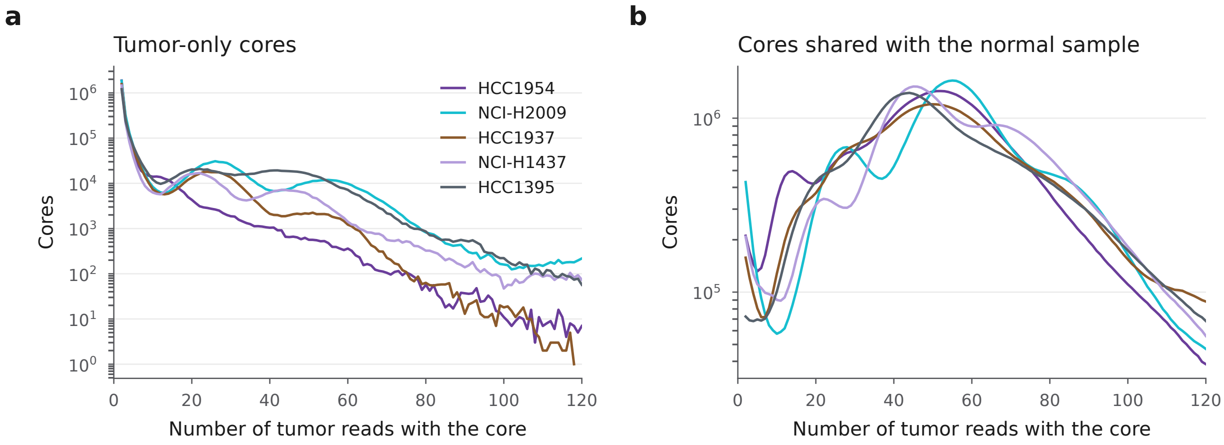

**Supplementary Figure S4:** Count spectra of the cores in the tumor samples of the CASTLE pairs at full depth: tumor-only cores (a) and cores shared with the normal sample (b), by the number of tumor reads that contain the core (at least two).

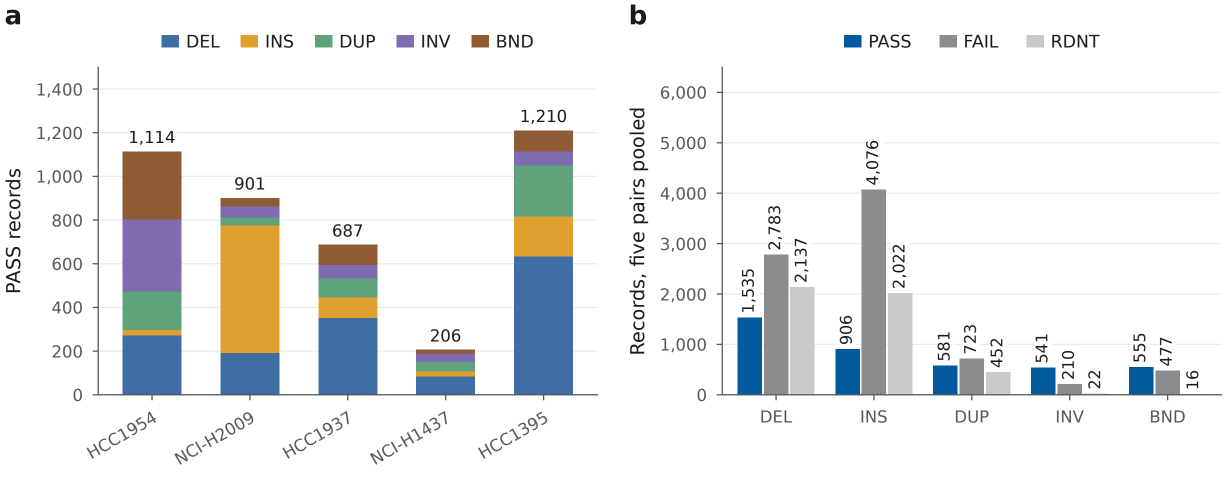

**Supplementary Figure S5:** Records of the five CASTLE pairs. (a) PASS records of each pair by SV type. (b) Records by SV type and by the filter assigned after classification, pairs pooled.

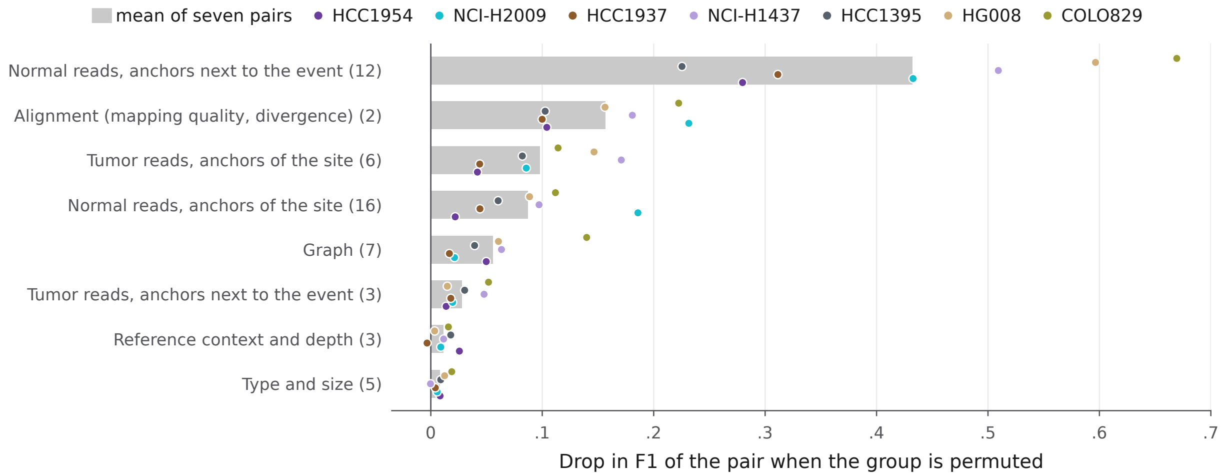

**Supplementary Figure S6:** Drop in F1 score (training labels) when the features of one group are permuted among the full-depth records of a pair, which is classified again with the same models. Bars show the mean over the seven pairs, dots the individual pairs, and parentheses the number of features. The minimum read support and the four depth and purity values are constant within a run and not shown.

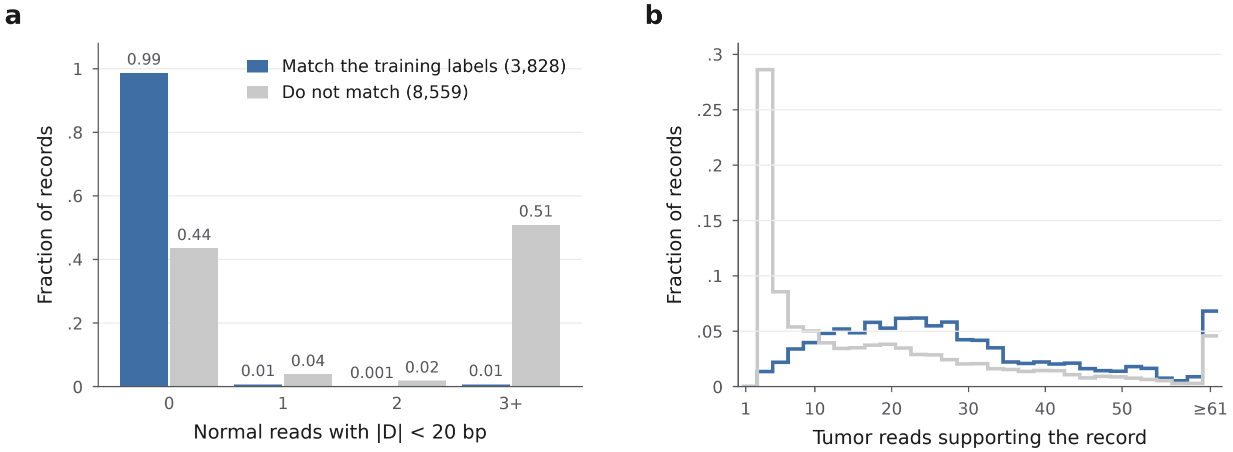

**Supplementary Figure S7:** Records of the five CASTLE pairs that match the training labels (3,828) and that do not (8,559), duplicates excluded. (a) Fraction of records by the number of normal reads with $|D| < 20\,\text{bp}$. (b) Fraction of records by the number of supporting tumor reads.

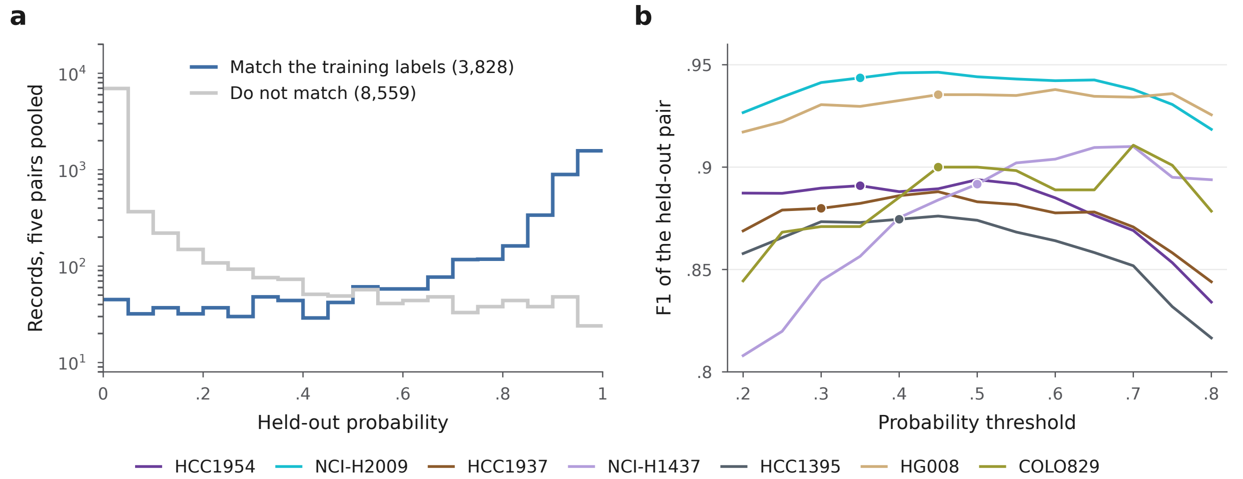

**Supplementary Figure S8:** (a) Held-out probabilities of the records of the five CASTLE pairs that match the training labels and that do not (duplicates excluded). (b) F1 score of each of the seven pairs on its training labels as a function of the threshold. The dot marks the threshold selected for the full-depth run.

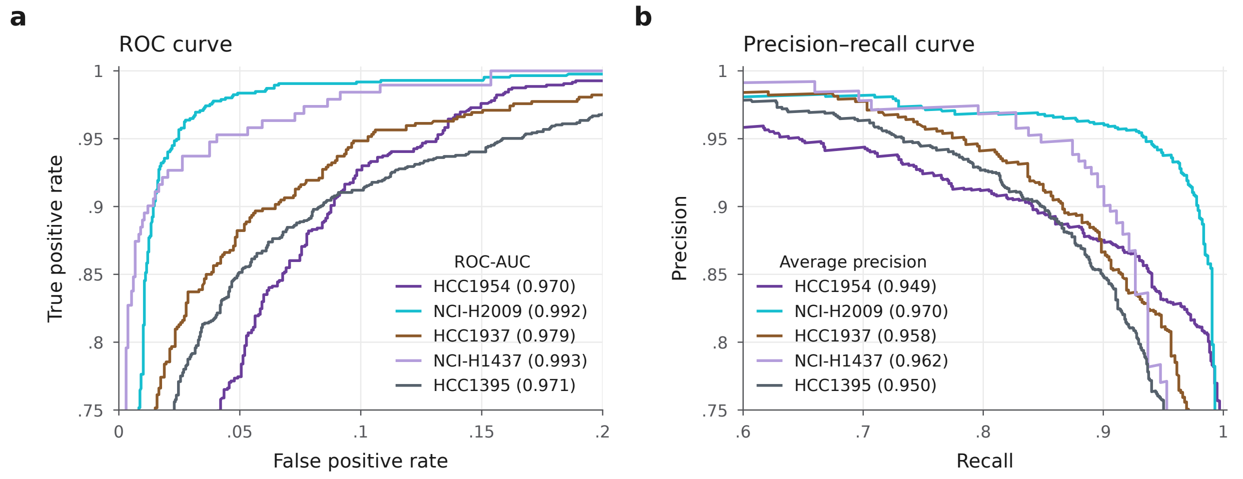

**Supplementary Figure S9:** Held-out probabilities of the records of the five CASTLE pairs at full depth, each pair classified by the model trained on the other four (RDNT records excluded, training labels): ROC curves (a) and precision–recall curves (b), with the ROC-AUC and the average precision in parentheses; axes zoomed to the curves.

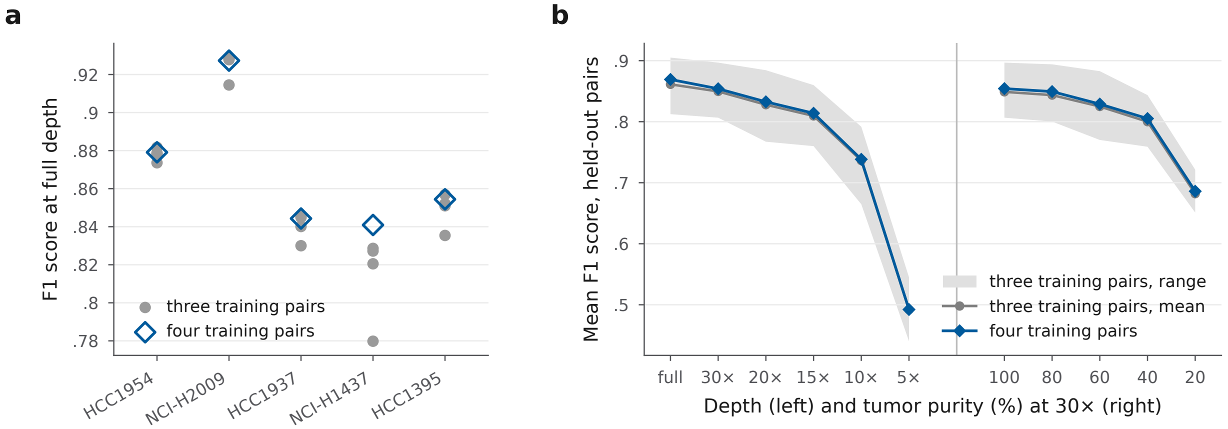

**Supplementary Figure S10:** Training on three instead of four CASTLE pairs (training labels). (a) F1 score of each pair at full depth with the four models trained on three of the other four pairs (gray) and with the model trained on the other four pairs (blue). (b) Mean F1 score of the held-out pairs at each depth and purity, with the mean and range over the 10 three-pair splits.

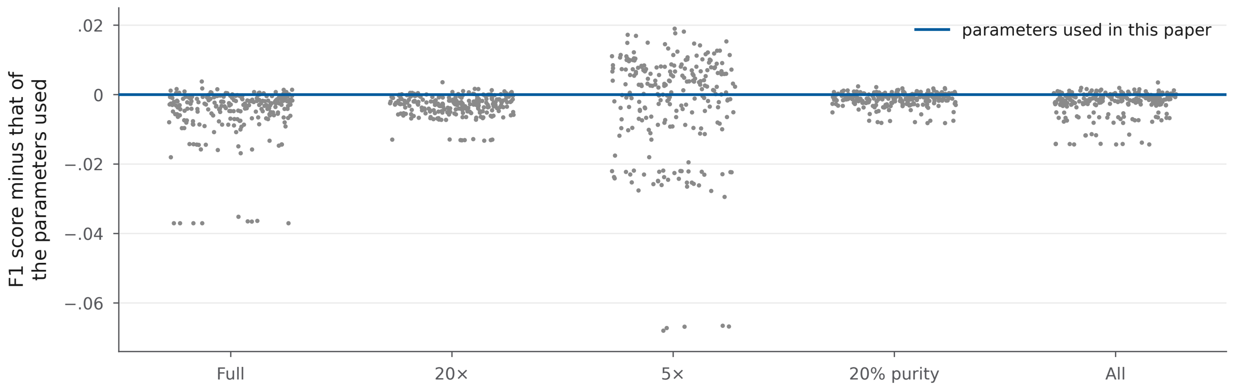

**Supplementary Figure S11:** F1 score of each of the 243 parameter sets minus that of the chosen parameters, as the mean of the five CASTLE pairs on the training labels at full depth, at 20×, at 5×, at 20% purity, and over all depths and purities except 100% purity (all).

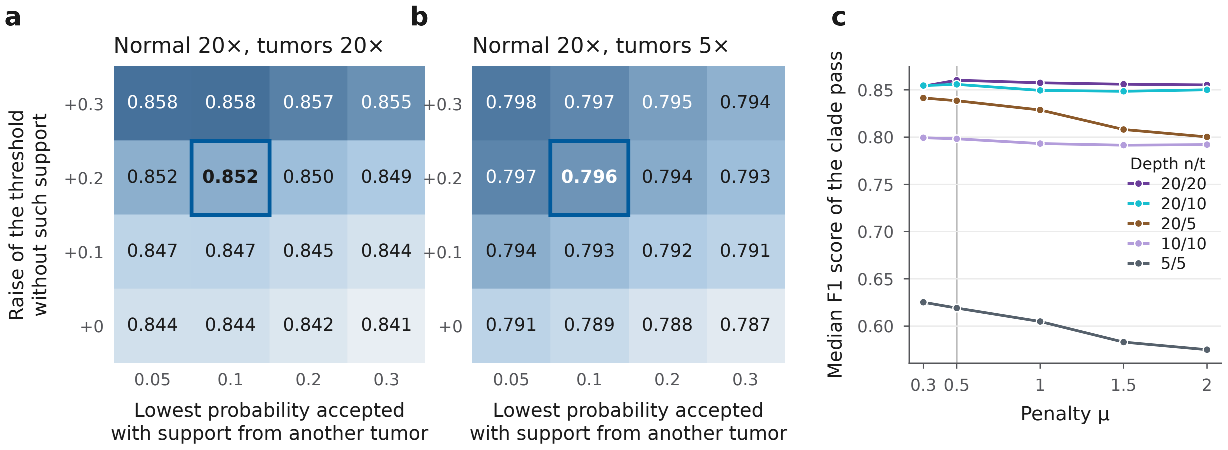

**Supplementary Figure S12:** Median F1 score of the 20 simulated tumors in the joint run. (a, b) First pass at 20/20 (a) and 20/5 (b) for different constants of the acceptance rule; rows give the margin above the threshold required without support from another tumor, and columns the lowest probability accepted with such support; the chosen values are outlined. (c) Clade pass for different values of the penalty $\mu$ of a tumor without a record, at the five depths; the chosen value is marked.

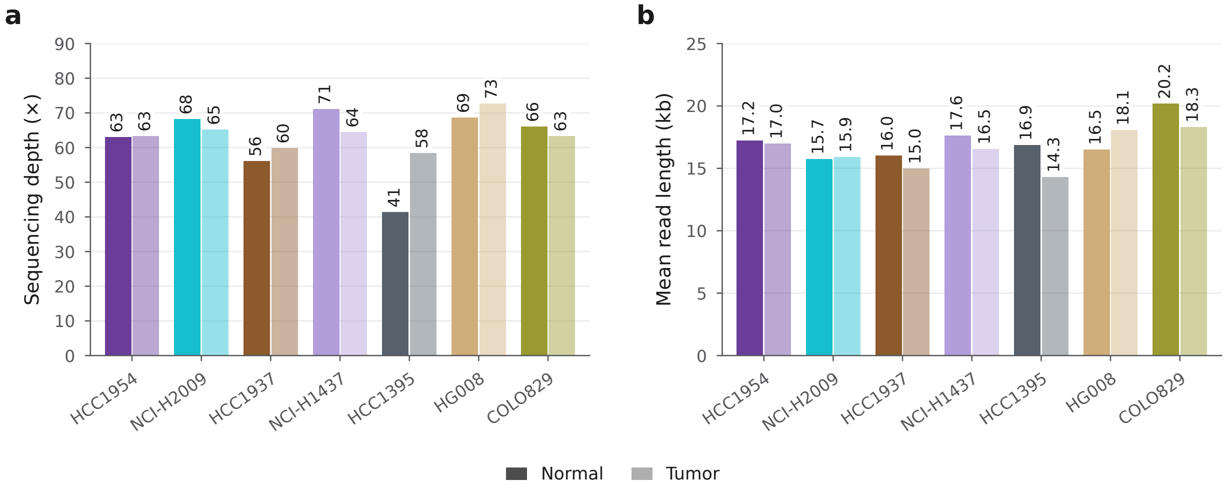

**Supplementary Figure S13:** The read sets of the seven tumor–normal pairs. (a) Sequencing depth (sequenced bases divided by 3.1 Gbp). (b) Mean read length.

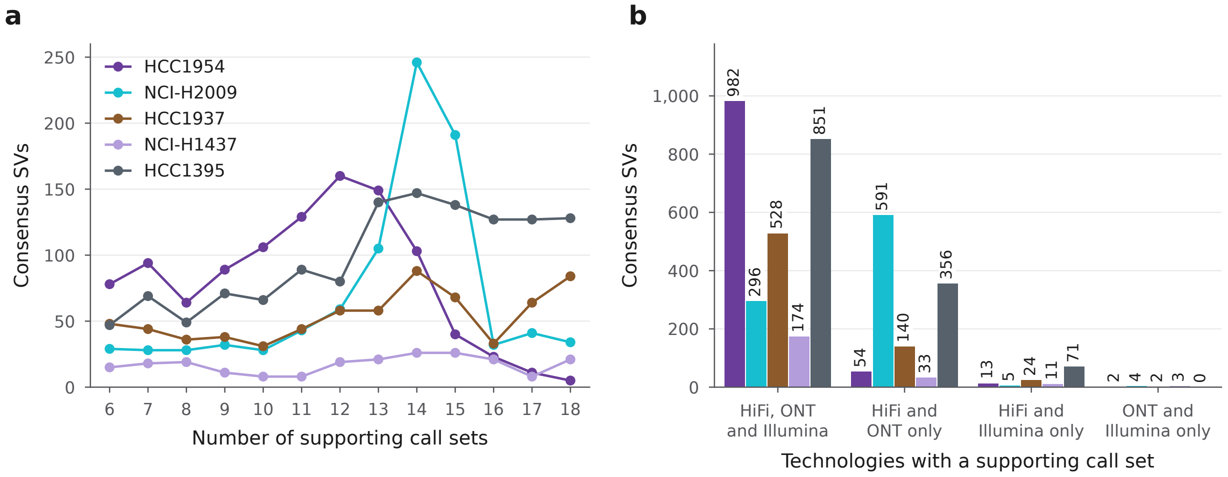

**Supplementary Figure S14:** Consensus of 18 call sets. (a) SVs by the number of call sets that contain them. (b) SVs by the technologies of these call sets, with the pair colors of (a).

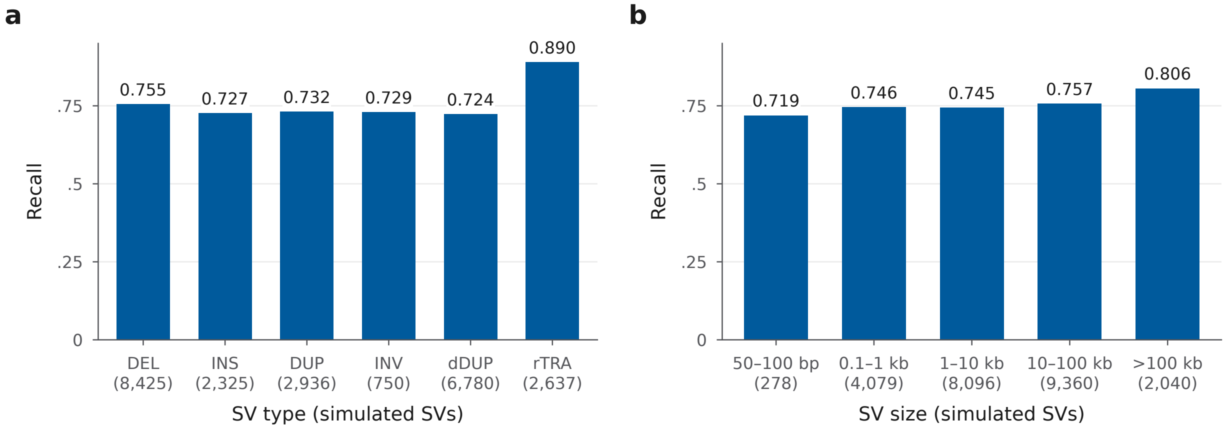

**Supplementary Figure S15:** Recall of the 20 pairwise runs on the simulated dataset at 20/20, tumors pooled, by SV type (a) and size (b). The number of simulated SVs, summed over the tumors, is given below each bar.

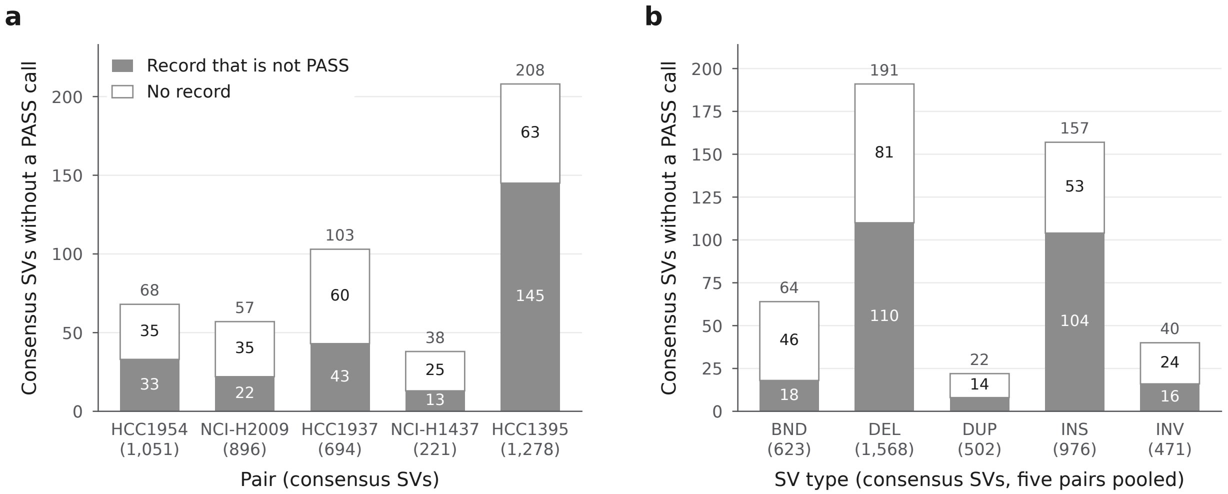

**Supplementary Figure S16:** Consensus SVs without a PASS call of JANUS (both breakpoints within 500 bp), split into SVs with a record that is not PASS and SVs without a record, by pair (a) and by SV type, pairs pooled, with inverted junctions counted as inversions (b); numbers in parentheses are the consensus SVs of the pair or type.

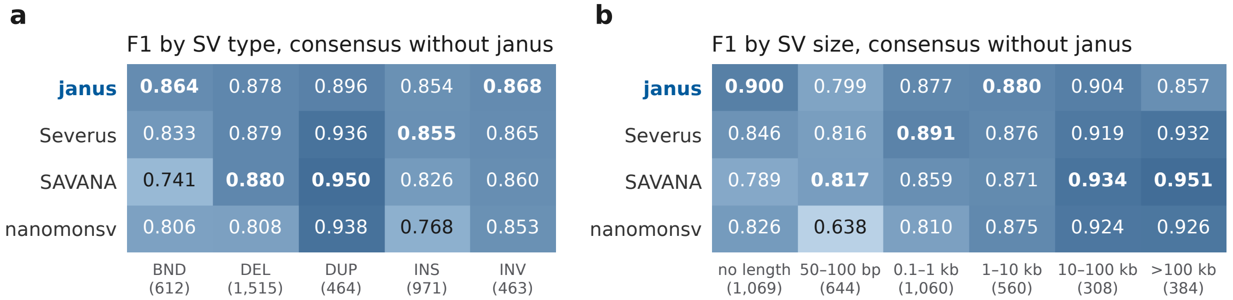

**Supplementary Figure S17:** F1 score by SV type (a) and size (b) against the consensus of the 17 call sets other than JANUS, five pairs pooled, with inverted junctions counted as inversions. The highest value of each column is in bold and the number of consensus SVs in parentheses.

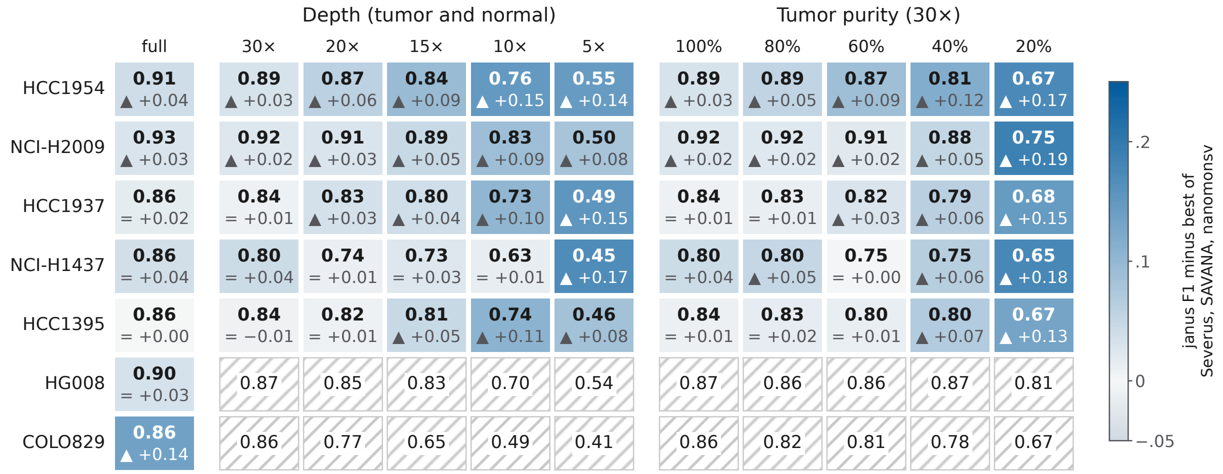

**Supplementary Figure S18:** F1 score of JANUS per pair and setting against the consensus of 18 call sets of the full-depth pair (HG008 and COLO829 against their truth sets), and below it, as color, its difference from the highest F1 score of Severus, SAVANA, and nanomonsv. Upward and downward triangles mark differences whose bootstrap 95% interval lies above or below zero, and = an interval that contains zero; the color scale runs from −0.05 to +0.25. Hatched cells have no runs of the other callers.

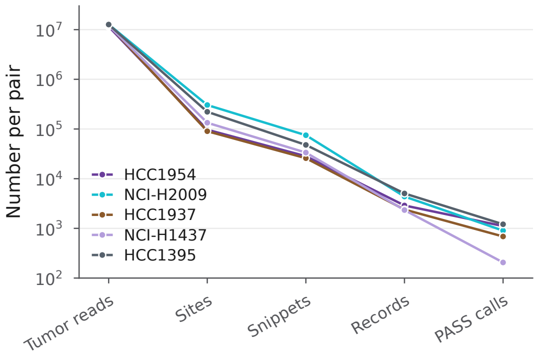

**Supplementary Figure S19:** Tumor reads, sites, snippets, records, and PASS calls per pair.
